# Fluorescence lifetime enables high-resolution analysis of neuromodulator dynamics across time and animals

**DOI:** 10.1101/2022.09.28.510014

**Authors:** Pingchuan Ma, Peter Chen, Elizabeth Tilden, Samarth Aggarwal, Anna Oldenborg, Yao Chen

**Affiliations:** Department of Neuroscience, Washington University in St. Louis, St. Louis, MO 63110; Ph.D. Program in Neuroscience, Washington University in St. Louis; Master’s Program in Biomedical Engineering, Washington University in St. Louis

**Keywords:** Optical sensors, neuromodulator, fluorescence lifetime, acetylcholine, sleep, running, behavior states, high-resolution dynamics, chronic, tonic

## Abstract

The dynamics of neuromodulators are essential for their functions. Optical sensors have transformed the study of neuromodulators because they capture neuromodulator dynamics with high spatial and temporal resolution. However, fluorescence intensity-based sensors are restricted to measure acute changes within one animal over a short period because intensity varies with sensor expression level and excitation light fluctuation. In contrast, fluorescence lifetime is impervious to sensor expression level or excitation light power, allowing comparison between individuals and across long periods. Here, we discover fluorescence lifetime response in multiple intensity-based neuromodulator sensors. Using the acetylcholine sensor GRAB_ACh3.0_ to investigate the power of lifetime measurement, we find that fluorescence lifetime correlates with animal behavior states accurately despite varying excitation power or changes in sensor expression level across weeks and animals. Thus, fluorescence lifetime of neuromodulator sensors enables comparison of neuromodulator dynamics at high resolution between animals and for chronic time scales.

## INTRODUCTION

Neuromodulators such as acetylcholine and dopamine can reconfigure neural circuits and transform animal behaviors^1–11^.They play important roles in normal physiology and their dysregulation is implicated in neurological and psychiatric disorders^12–19^. Despite decades of research on neuromodulators, many questions remain. Notably, tonic and phasic firing of neuromodulator-releasing neurons result in distinct changes in synaptic properties and behavior^20–24^, but we know very little about when tonic versus phasic changes of neuromodulators occur during animal behavior. In addition, neuromodulators are released widely into many brain regions^25^, but it is unclear whether their release is differentially regulated in different regions. Finally, most drugs for psychiatric disorders target neuromodulators or their receptors^13, 16, 17, 26–29^, but we cannot easily compare neuromodulator levels between control and disease models, between pre-drug and post-drug periods, and we understand even less whether these drugs alter transient or sustained levels of neuromodulators. Thus, to advance our understanding of the function of neuromodulators in animal behavior, we need methods to capture both transient and sustained neuromodulator changes, and to compare these changes between brain regions, between disease models, and across chronic periods.

Current methods to analyze neuromodulators have provided important information on their involvement in behavior but do not allow the dissection between transient and sustained neuromodulator changes. Classical methods such as microdialysis and electrochemical methods allow comparison of neuromodulator concentration over long periods of time and between animals^30–34^. However, these methods lack spatial resolution, temporal resolution, or chemical specificity. Genetically encoded optical reporters of neuromodulators are now transforming the field of neuromodulation due to their high spatial and temporal resolution^35–39^. Most of these optical sensors are derived from the membrane receptors for the specific neuromodulators, and they increase in fluorescence intensity upon ligand binding. However, fluorescence intensity does not only respond to changing neuromodulator concentrations, but also varies with excitation light power and sensor expression level, which occur across long time periods, between brain regions, and between animals (Fig. 8). As a result, intensity measurement cannot be used to compare neuromodulator concentrations across these domains, or to quantitate changes in tonic levels of neuromodulators (Fig. 8). Therefore, an ideal sensor would combine the benefits of classical methods and fluorescence intensity-based sensors to allow high-resolution measurement of neuromodulator concentrations across time and animals.

Fluorescence lifetime imaging microscopy (FLIM) measurement of optical sensors could fulfil the requirement of such an ideal sensor. Fluorescence lifetime measures the time between excitation and light emission of a fluorophore and is therefore independent of sensor expression levels or fluctuation in excitation light power^38, 40–43^. FLIM has been frequently used to track the conformational change of biosensors and has been used successfully to uncover spatiotemporal dynamics of intracellular signals and voltage. Whereas most FLIM sensors involve dyes or are based on Förster Resonance Energy Transfer^41, 44–51^, most neuromodulator sensors are single fluorophore-based. Although a few single-fluorophore protein-based sensors show fluorescence lifetime change^52–54^, the majority of them do not, and it is hard to predict whether a given sensor will show fluorescence lifetime change. Importantly, no genetically encoded neuromodulator sensor has been reported to show a lifetime change. Furthermore, FLIM is rarely used to make comparison across animals or chronic time periods in vivo. Thus, it is unclear whether any intensity-based neuromodulator sensors can display a fluorescence lifetime change; nor is it known whether FLIM is a viable technique to predict neuromodulator levels across excitation light powers, different individual animals, and chronic time periods.

Here, we tested whether any existing neuromodulator sensors^55–60^ showed a fluorescence lifetime change and discovered lifetime response in multiple sensors. We used the acetylcholine (ACh) sensor GRAB_ACh3.0_ (GPCR-Activation-Based acetylcholine sensor 3.0)^56^ to investigate the power of lifetime measurement because it displayed the largest dynamic range. Like intensity, FLIM measurement of GRAB_ACh3.0_ can detect transient ACh changes, is dose sensitive, and shows high spatial and temporal resolution. In contrast to intensity, FLIM measurement correlates much better with ACh-associated behavior states, despite laser power fluctuation or sensor expression change across weeks or animals. Our results have broad implications beyond ACh sensors. Methodologically, these results demonstrate the power of FLIM for neuromodulator measurement, highlighting the importance to convert many existing fluorescence intensity-based neuromodulator sensors into lifetime-based sensors. Biologically, FLIM measurement of neuromodulator sensors enables us to simultaneously capture both transient and sustained changes of neuromodulators, promising to disambiguate phasic and tonic contributions across animals, disease models, brain regions, and over long periods of time.

## RESULTS

### Fluorescence Lifetime Responses of Neuromodulator Sensors

We tested whether any existing intensity-based neuromodulator sensors also showed a fluorescence lifetime change (Fig. 1A). We expressed individual sensors in human embryonic kidney (HEK) 293T cells and measured sensor fluorescence intensity and lifetime with two-photon fluorescence lifetime imaging microscopy (2pFLIM). Surprisingly, in addition to fluorescence intensity changes (Fig. 1B; GRAB_ACh3.0_^56^, n = 18, p < 0.0001; intensity-based ACh sensing fluorescent reporter (iAChSnFR)^58^, n = 11, p = 0.001; 5-HT sensor GRAB_5-HT_^57^, n = 29, p < 0.0001; norepinephrine (NE) sensor GRAB_NE_^60^, n = 15, p < 0.0001; dopamine (DA) sensor GRAB_DA2m_^59^, n = 19, p < 0.0001), multiple sensors showed a significant fluorescence lifetime change in response to saturating concentrations of the corresponding neuromodulators (Fig. 1B; GRAB_ACh3.0_, p < 0.0001; iAChSnFR, p = 0.001; GRAB_5-HT_, p = 0.0004; GRAB_NE_, p= 0.1514; GRAB_DA2m_, p = 0.001). These results demonstrate that single fluorophore-based neuromodulator sensors can show fluorescence lifetime responses.

**Figure 1.**
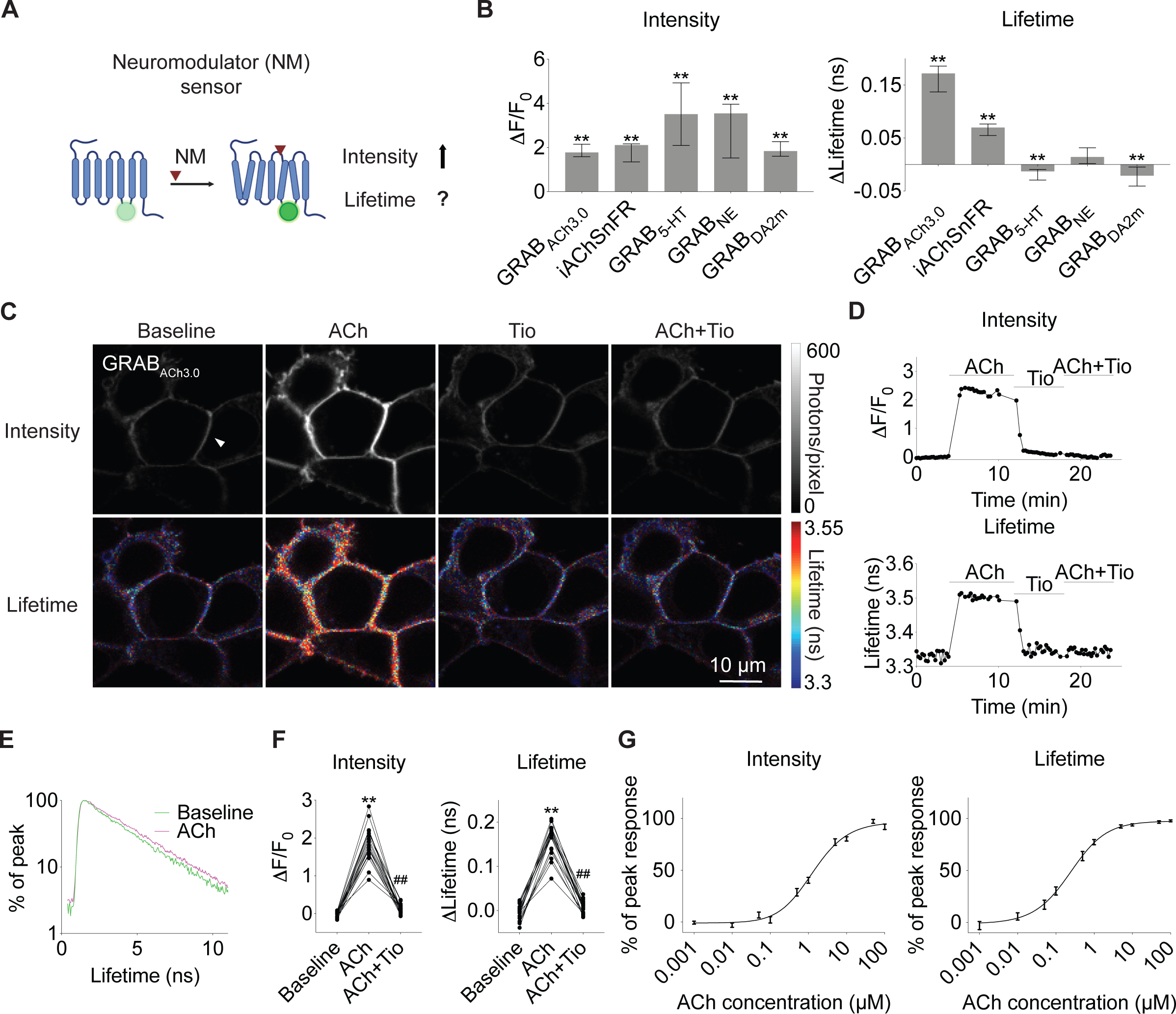
The ACh sensor GRAB_ACh3.0_ shows fluorescence lifetime response. **(A)** Schematic illustrating the question under investigation: neuromodulator sensors show fluorescence intensity increase, but it is unclear whether they show any fluorescence lifetime change. **(B)** Summaries of fluorescence intensity and lifetime changes of different neuromodulator sensors in response to saturating concentrations of the corresponding neuromodulators in HEK 293T cells. Wilcoxon test, **p < 0.01, vs baseline change. Data are represented as median with interquartile range. **(C-D)** Representative heatmaps (C) and traces (D) showing fluorescence intensity (upper panels) or fluorescence lifetime (lower panels) of GRAB_ACh3.0_ in response to saturating concentration of ACh (100 μM) with the cholinesterase inhibitor (AChEi) Donepezil (5 μM), mAChR antagonist Tiotropium (Tio, 5 μM), or ACh+Tio+Don in HEK 293T cells. The traces in D are from the cell denoted by a triangle in C. **(E)** Histogram of fluorescence lifetime of GRAB_ACh3.0_ sensor under baseline and with 100 μM ACh. **(F)** Summaries of intensity and fluorescence lifetime changes of GRAB_ACh3.0_ sensor in HEK 293T cells. Note these data are the same as those displayed for GRAB_ACh3.0_ in Fig. 1B. Friedman one-way ANOVA test with Dunn’s multiple comparison, **p < 0.01 vs baseline, ^##^p < 0.01 vs ACh. **(G)** Summaries of the dose-dependent intensity and fluorescence lifetime change of GRAB_ACh3.0_ sensor in response to different concentrations of ACh in the presence of 5 μM AChEi Donepezil. Data are represented as mean with standard error of the mean (SEM). See also Figure S1.

We subsequently used the ACh sensor GRAB_ACh3.0_^56^ to investigate the power of lifetime measurement because of the following reasons. First, ACh is one of the best-characterized neuromodulators. It increases during defined behavior state transitions, such as from resting to running^56, 61–63^, and from non-rapid eye movement (NREM) sleep to REM sleep^56, 64–69^, thus making it feasible to test the power of the technology with known ground truth. Second, ACh is one of the most important neuromodulators in the brain^17, 70^, playing critical roles in neuronal processes including learning and memory^71^, attention^72^, and sleep^73^. Third, GRAB_ACh3.0_ showed the largest fluorescence lifetime change among all the neuromodulator sensors tested (Fig. 1B; median of 0.17 ns with interquartile range of 0.14-0.19 ns in response to 100 µM ACh; n = 18; p < 0.0001). The large dynamic range makes it easier to explore the power of lifetime measurement in vivo. In the initial characterization of GRAB_ACh3.0_, like intensity, lifetime of GRAB_ACh3.0_ increased in response to saturating concentration of ACh (100 μM) and this increase was blocked by the addition of the muscarinic ACh receptor (mAChR) antagonist tiotropium (Tio, 5 μM) (Fig. 1C-1D, 1F; n = 18; adjusted p = 0.0007 for intensity and < 0.0001 for lifetime; ACh+Tio vs ACh). Furthermore, a mutant sensor that does not bind ACh (GRAB_ACh3.0mut_) did not show any intensity or fluorescence lifetime change in response to ACh (Fig. S1; n = 5; p = 0.31 for intensity and 0.63 for lifetime). Importantly, the fluorescence lifetime histogram of GRAB_ACh3.0_ showed slower decay with 100 μM ACh than without ACh at baseline (Fig. 1E), indicating that ACh binding increases fluorescence lifetime. Thus, both intensity and lifetime respond to ACh in cells expressing GRAB_ACh3.0_.

To test whether lifetime of GRAB_ACh3.0_ responds to graded ACh, we measured the dose response curve of GRAB_ACh3.0_. In response to different concentrations of ACh ranging from physiologically relevant to saturating concentrations (1 nM to 100 μM)^74–76^, fluorescence lifetime of GRAB_ACh3.0_ in HEK cells showed a dose-dependent increase (Fig. 1G; n = 13). These results indicate that lifetime measurement of GRAB_ACh3.0_ report graded ACh increase.

In principle, an increase in fluorescence lifetime of cells expressing GRAB_ACh3.0_ could be due to true lifetime response to ACh by GRAB_ACh3.0_, or due to an increase in intensity of GRAB_ACh3.0_ relative to the autofluorescence of cells without any change of GRAB_ACh3.0_ lifetime. The latter possibility exists because both the fluorescent sensor and autofluorescence contribute to fluorescence measurement of cells, and the lifetime of GRAB_ACh3.0_ is longer than that of autofluorescence (Fig. S2A). To test the null hypothesis that GRAB_ACh3.0_ showed no lifetime change, we performed computational simulations to test how much cellular lifetime would increase if GRAB_ACh3.0_ only increased in intensity and not lifetime. For the simulation, we constructed photon populations of GRAB_ACh3.0_ sensor as double exponential decay (Fig. S2B). Subsequently, we sampled from this population with low and high photon numbers corresponding to measurements at 0 and 100 µM ACh respectively (Fig. 2A). We additionally added autofluorescence based on measurement in cells without sensor expression. Our simulation showed that if the sensor itself did not show any fluorescence lifetime increase, an increase in intensity only caused a small increase of overall lifetime (Fig. 2B; from 3.242 ± 0.012 ns to 3.247 ± 0.0065 ns; n = 500 simulations for both low and high photons). In contrast, the experimentally measured lifetime increase in response to 100 µM ACh was much larger (Fig. 2B; n = 3; mean difference = 0.19 ns), more than 10 times of the standard deviation (0.014 ns) of the difference between low and high photons from simulation. Therefore, the observed fluorescence lifetime response in cells expressing GRAB_ACh3.0_ is not solely due to an increase in fluorescence intensity. Rather, GRAB_ACh3.0_ sensor itself responds to ACh with authentic fluorescence lifetime increase.

**Figure 2.**
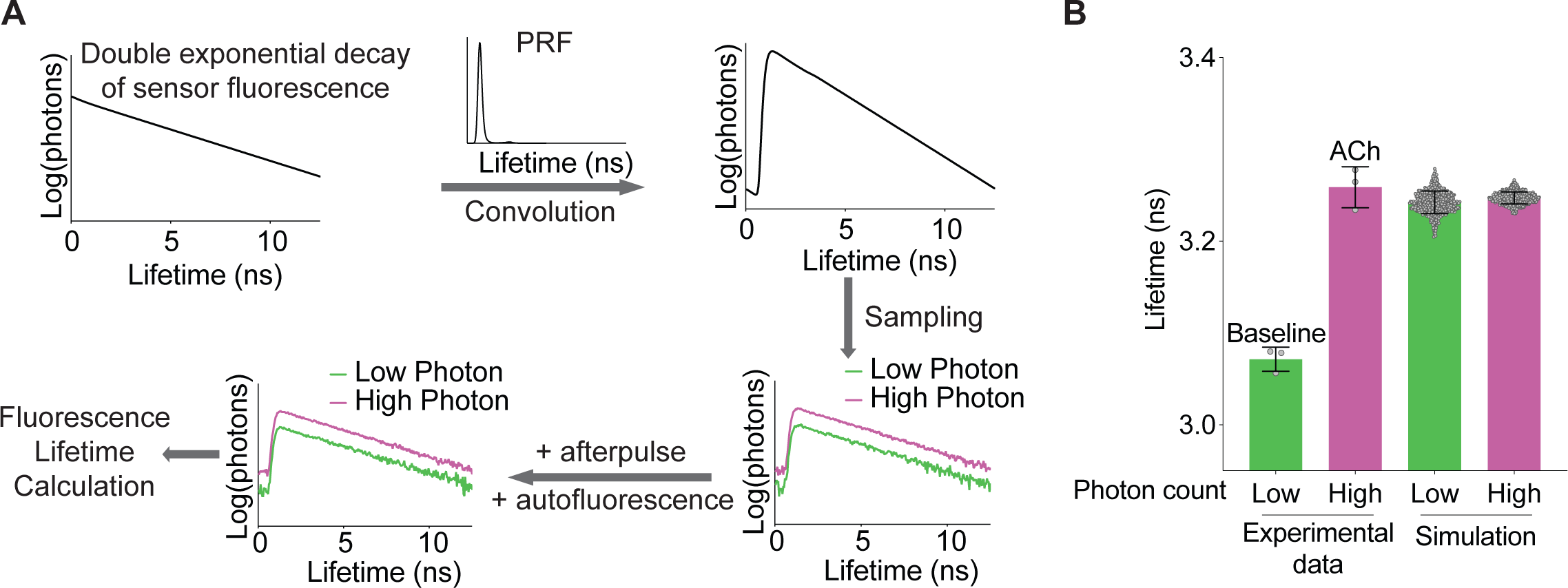
Simulation reveals authentic fluorescence lifetime response of GRAB_ACh3.0_. **(A)** Schematic illustrating the process of simulation. Fluorescence lifetime histogram of the sensor is modeled as a double exponential decay, convolved with measured pulse response function (PRF), and sampled with different number of photons. Subsequently, afterpulse and autofluorescence (sampled from measured distribution) are added. Empirical fluorescence lifetime was then calculated from the simulated distribution. **(B)** Fluorescence lifetime distribution of cells expressing GRAB_ACh3.0_ based on experimental data (n = 3) and based on simulation (n = 500 simulations under each condition). Experimental data were collected in the absence or presence of ACh (100 μM). Simulation assumed only intensity change, and no lifetime change of the fluorescence sensor, and simulated with low or high photon counts corresponding to baseline and ACh conditions respectively. Data are represented as mean with standard deviation. See also Figure S2.

### Fluorescence lifetime of ACh sensor detects transient ACh change in the brain

To test whether fluorescence lifetime of GRAB_ACh3.0_ can report ACh levels in brain tissue, we delivered the reporter via adeno-associated virus (AAV) injection to CA1 pyramidal neurons of the mouse hippocampus (Fig. 3A). Bath application of ACh (1 μM and 100 μM) induced both fluorescence lifetime (Fig. 3B-3C; n = 8; adjusted p = 0.023 for baseline vs 1 μM, baseline vs 100 μM, and 1 μM vs 100 μM) and intensity (Fig. S3A-S3B; n = 8; adjusted p = 0.023 for baseline vs 1 μM, baseline vs 100 μM, and 1 μM vs 100 μM) increase of GRAB_ACh3.0_. These results indicate that fluorescence lifetime of GRAB_ACh3.0_ is sensitive enough to report ACh increase in brain tissue.

**Figure 3.**
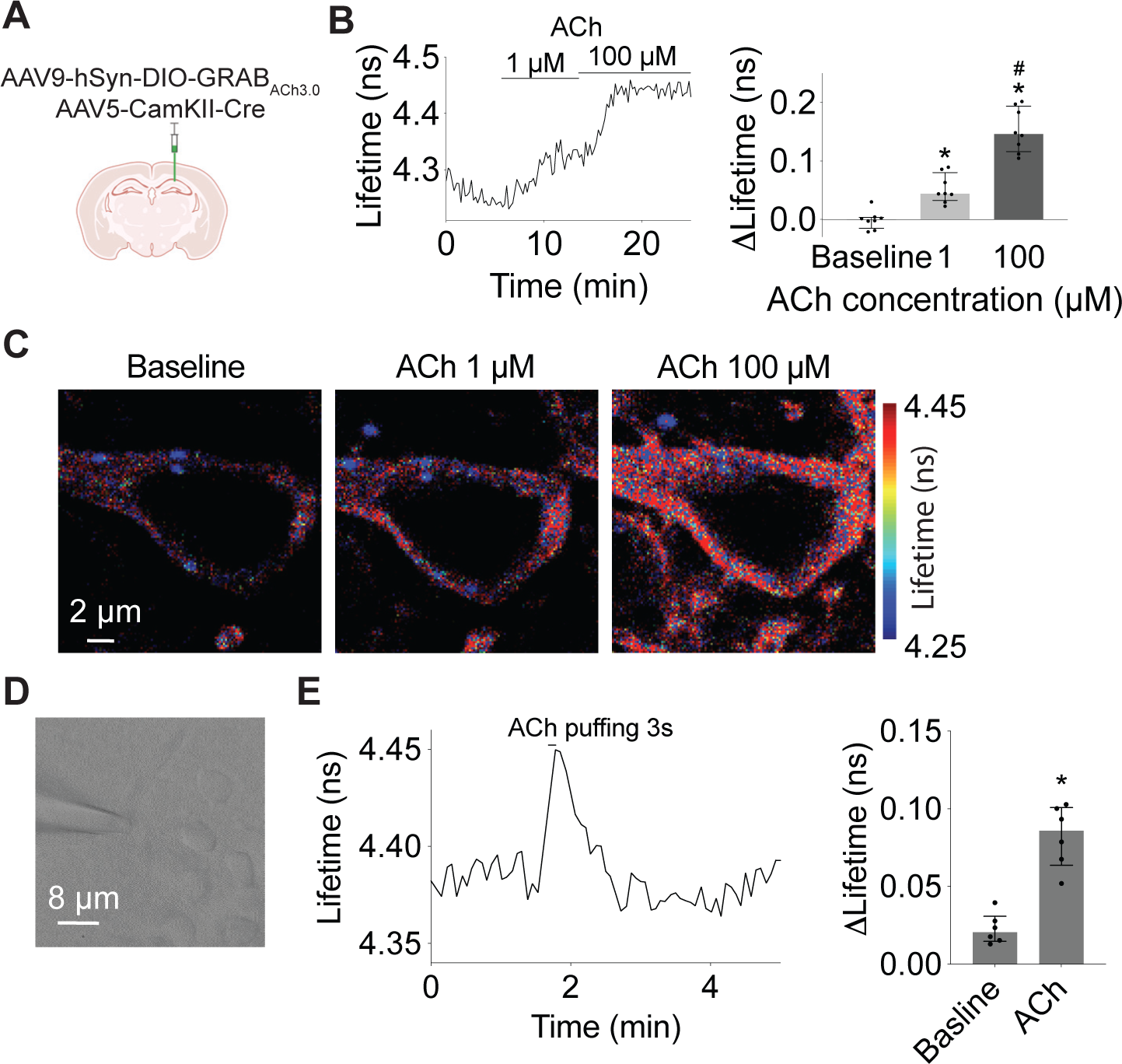
Fluorescence lifetime of GRAB_ACh3.0_ responds to transient ACh in brain tissue. **(A)** Illustration of expression of GRAB_ACh3.0_ in CA1 cells of hippocampus by AAVs. AAVs carrying Cre recombinase driven by the neuronal specific CamKII promoter and Cre-dependent GRAB_ACh3.0_ were delivered to CA1 region in the hippocampus of wild-type mice. **(B-C)** Example trace and summaries (B), as well as heatmaps (C) showing fluorescence lifetime of hippocampal CA1 pyramidal neurons expressing GRAB_ACh3.0_ in response to ACh (1 µM and 100 µM, with 5 µM AChEi Donepezil). Wilcoxon test with Bonferroni correction, *p < 0.05 vs baseline, ^#^p < 0.05 vs 1 µM. **(D)** Gradient contrast image showing puffing of ACh onto a CA1 pyramidal neuron with a glass pipette connected to a Picospritzer. **(E)** Example trace and summaries showing fluorescence lifetime of GRAB_ACh3.0_ in CA1 pyramidal neurons in response to a 3-second puff of ACh (200 μM). Wilcoxon test, *p < 0.05 vs baseline. Data in B and E are represented as median with interquartile range. See also Figure S3.

For fluorescence lifetime measurement of GRAB_ACh3.0_ to be useful in biological applications, it needs to be sensitive enough to detect transient ACh in the brain. To test this, we puffed ACh (200 μM) onto the soma of CA1 pyramidal neurons (Fig. 3D) at temporal duration (3 seconds) comparable to ACh release measured in behaving animals in vivo^77^. Both fluorescence lifetime (Fig. 3E; n = 6; p = 0.031) and intensity (Fig. S3C; n = 6; p = 0.031) of GRAB_ACh3.0_ increased in response to ACh delivery, indicating that lifetime of GRAB_ACh3.0_ can report in brain tissue ACh release that is temporally relevant and transient.

Together, these results show that like intensity, fluorescence lifetime of GRAB_ACh3.0_ can report transient increase of ACh in the brain.

### Fluorescence lifetime of ACh sensor is independent of laser power

Unlike intensity, fluorescence lifetime should be independent of laser power fluctuation. To explore the extent of this advantage, we measured both fluorescence lifetime and intensity under different laser excitation powers. As expected, fluorescence intensity of GRAB_ACh3.0_ increased with increasing laser power (Fig. 4A-4B; n = 10; adjusted p = 0.0005 for baseline and < 0.0001 for ACh, low vs high laser power). Both laser power and the presence of ACh contributed significantly to the variability of fluorescence intensity across cells (Fig 4C, p < 0.0001 for both ACh and laser power). Only 49% of sensor intensity variance could be explained by ACh concentrations (Fig. 4C). In contrast, fluorescence lifetime of the ACh sensor was stable across different laser powers (Fig. 4A-4B; n = 10; adjusted p = 0.71 for baseline and 0.68 for ACh, low vs high laser power). Only the presence or absence of ACh, and not laser power, significantly contributed to the variation of fluorescence lifetime across cells (Fig 4C, p < 0.0001 for ACh, p = 0.18 for laser power). Notably, the majority (73%) of the variance of sensor lifetime could be explained by ACh concentration, with minimal contributions from laser power (0.11%) or cell identity (23%; Fig. 4C). Together, these results indicate that fluorescence lifetime is a more reliable measurement of ACh concentration than fluorescence intensity under fluctuating laser powers.

**Figure 4.**
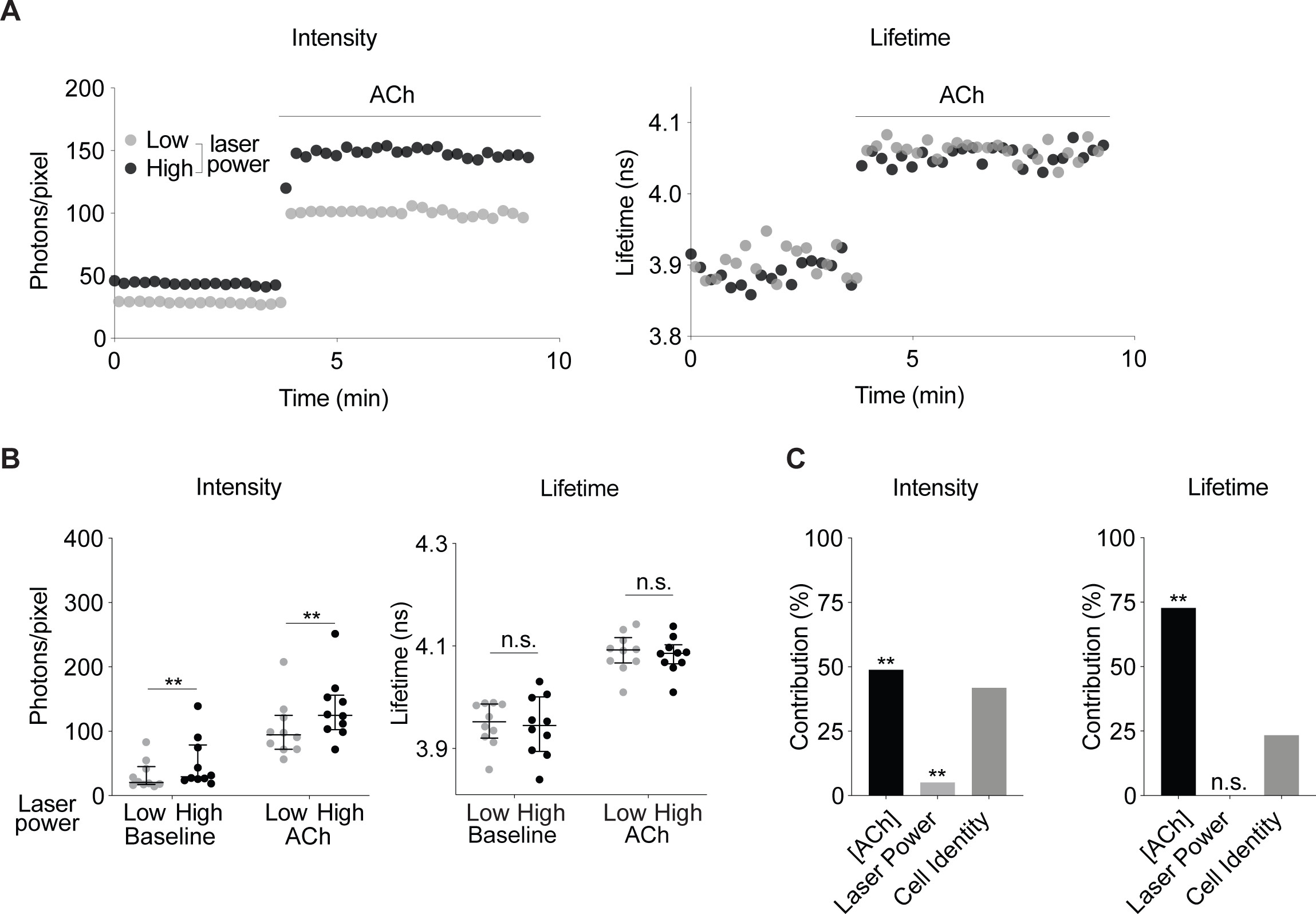
Fluorescence lifetime is stable across different excitation light powers. **(A)** Representative traces of intensity (left) and fluorescence lifetime (right) of HEK 293T cells expressing GRAB_ACh3.0_ in response to ACh (100 μM, with 5 µM AChEi Donepezil), imaged at different laser powers. **(B)** Summaries of intensity and fluorescence lifetime of cells expressing GRAB_ACh3.0_ under different laser powers, and in the absence and presence of ACh. Two-way ANOVA with Šídák’s multiple comparison, **p < 0.01, n.s. not significant, low vs high laser power. Data are represented as median with interquartile range. **(C)** Two-way ANOVA analysis showing the contribution to the total variance of the measurements due to ACh concentration, laser power, or cell identities. **p < 0.01, n.s. not significant. See also Figure S4.

### Fluorescence lifetime is consistent within a cell and between cells

If absolute fluorescence lifetime were to be used to predict ACh concentrations, lifetime values would need to be stable within a cell for a given ACh concentration, and consistent between cells. To test the stability of lifetime within a cell, we repeatedly applied ACh (1 µM). Like intensity, fluorescence lifetime was consistent within a cell across repeated application of the same concentration of ACh (Fig. S4A-S4B; n = 8; p > 0.99 for intensity and p = 0.95 for lifetime, 1^st^ vs 2^nd^ flow-in). Thus, lifetime is consistent for a given ACh concentration within a cell.

To test whether absolute fluorescence lifetime correlates well with ACh concentration between cells, we measured both lifetime and intensity exposed to a specified ACh concentration that is comparable to that reported in vivo^74–76^. As expected, fluorescence intensity varied greatly between cells at a given ACh concentration (Fig. 5; 1 µM: coefficient of variation (CV) = 53.23% at baseline and 44.36% with ACh, n = 77 and 99; 10 µM: CV = 59.06% at baseline and 52.51% with ACh, n = 35 and 114), likely due to different sensor expression levels across cells. Although fluorescence intensity increased in response to ACh (Fig. 5; p<0.0001 for baseline vs ACh, both 1 µM and 10 µM ACh), intensity alone correlated poorly with ACh concentration (Fig. 5; baseline versus ACh, pseudo R^2^ = 0.12 for 1 µM ACh and 0.13 for 10 µM ACh). In contrast, for fluorescence lifetime, variation between cells was much smaller (Fig. 5; 1 µM: CV = 0.91% at baseline and 1.17% with ACh, n = 77 and 99; 10 µM: CV = 0.63% at baseline and 0.75% with ACh, n = 35 and 114). The signal-to-noise ratio was high. Absolute lifetime values correlated with ACh concentration with high accuracy (Fig. 5; baseline versus ACh, pseudo R^2^ = 0.77 for 1 µM ACh and 1 for 10 µM ACh). The variation of lifetime across cells was not due to the presence of varied amount of ACh at baseline (Fig. S5A; n = 13; p = 0.64 for baseline vs Tio), or varied amount of cholinesterase activity (Fig. S5B; p = 0.67; CV = 1.12% without and 1.01% with cholinesterase inhibitor (AChEi) Donepezil (5 μM); n = 40 and 61 respectively). In fact, the variability was comparable to the mutant sensor GRAB_ACh3.0mut_ that cannot bind ACh (Fig. S5C; p = 0.6041; CV s= 0.79% without and 0.92% with ACh; n = 42 and 53 respectively). These data suggest that lifetime variability between cells is likely due to the flexibility of sensor conformation. Furthermore, fluorescence lifetime, unlike fluorescence intensity, correlates with ACh concentration with high accuracy despite different sensor expression levels across individual cells.

**Figure 5.**
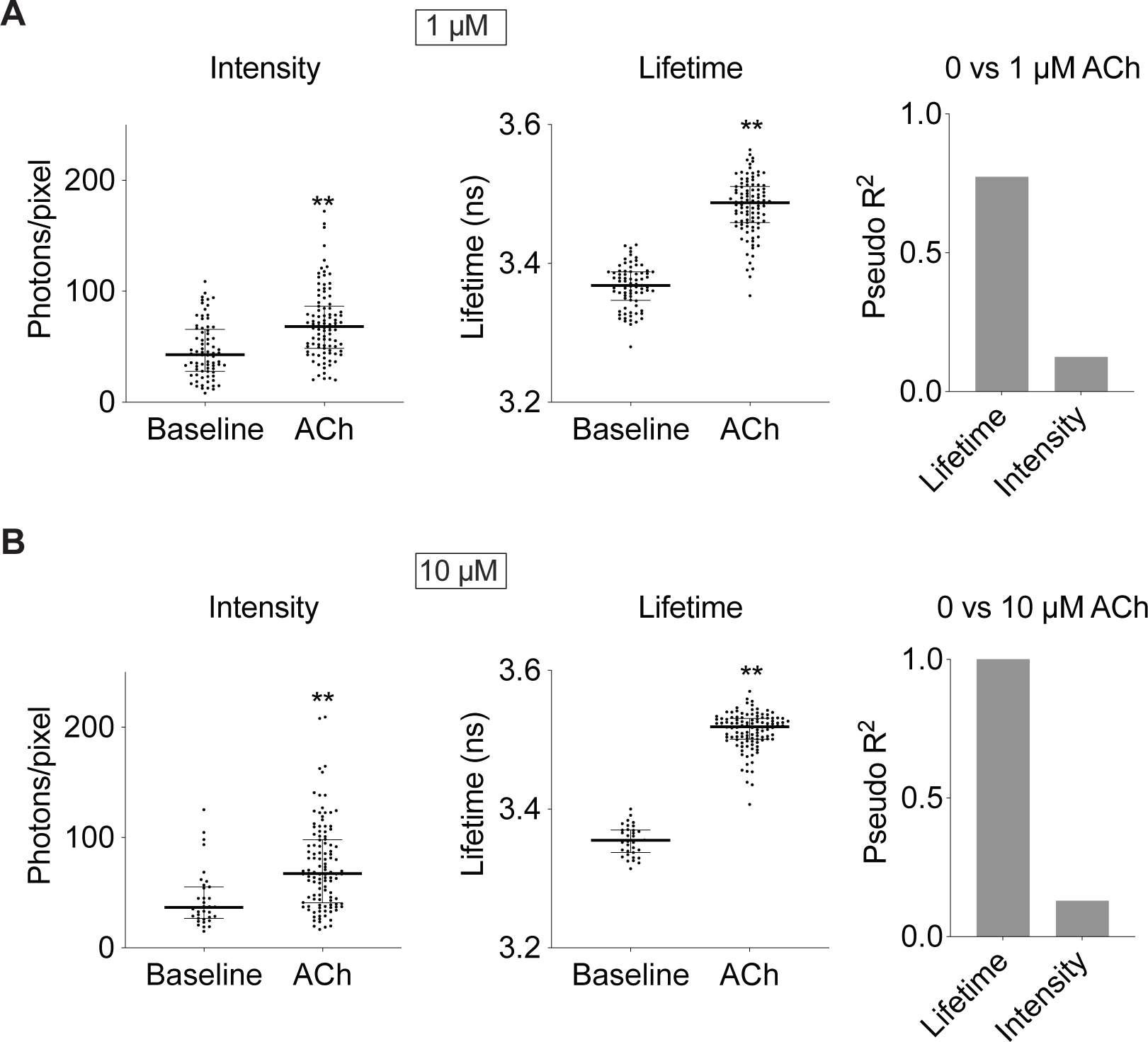
Fluorescence lifetime shows much less variability across cells and correlates better with ACh concentration than intensity. **(A-B)** Left: Distribution of intensity and fluorescence lifetime measurements of GRAB_ACh3.0_ in HEK 293T cells, at baseline and with different concentrations of ACh (1 μM and 10 μM, with 5 μM AChEi Donepezil). Mann-Whitney test, **p < 0.01 vs baseline. Data are represented as median with interquartile range. Right: Pseudo R^2^ values between intensity/lifetime and ACh concentrations based on logistic regression, showing lifetime measurement has much greater explanatory power than intensity for ACh concentration. See also Figure S5.

### Fluorescence lifetime correlates with ACh-associated running-resting states with high accuracy across individual mice and varying laser powers

Our goal is to compare ACh levels across imaging conditions, between mice, and chronic time scales such as weeks or months at high temporal resolution. We thus tested whether lifetime can measure both acute and sustained changes of neuromodulator concentrations in vivo, thus offering advantages of both intensity-based measurement of optical sensors and microdialysis. To assess lifetime measurement of acute changes, we tested whether lifetime of GRAB_ACh3.0_, like intensity, reports fast behavior state transitions correlated with ACh concentrations. To assess the potential of lifetime measurement to capture sustained changes, we used known ACh-correlated behavior states as ground truth, and asked whether lifetime measurement can accurately explain the variation of these behavior states across different laser powers, different individual mice, and different sensor expression levels across weeks.

Our proof-of principle experiments involve fluorescence lifetime photometry (FLiP) to measure lifetime and intensity simultaneously as mice transition between resting/running and different stages of sleep/wake. FLiP measures the bulk fluorescence from a population of cells surrounding the tip of the fiber implant, allowing for the measurement of neuromodulator dynamics in genetically defined neurons in a brain region in vivo^78^. The signal-to-noise ratio for the bulk signal is thus even higher than methods with cellular resolution. In fact, the variance of the signal is inversely proportional to the number of cells. Thus, if the bulk signal of ∼1000 cells were analyzed, the standard deviation of lifetime distribution would be 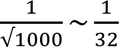 of the standard deviation across single cells (Fig. S6A), making FLiP a superb method to measure ACh level in vivo.

First, we tested whether fluorescence lifetime measurement of the ACh sensor increased as mice transitioned from resting to running, since ACh is high during running than resting^56, 61–63^. AAV virus carrying Cre-dependent GRAB_ACh3.0_ was delivered to hippocampal CA1 region of Emx1^IRES*cre*^ mice^79^, labelling excitatory neurons and a subset of glia with the ACh sensor (Fig. 6A). We recorded fluorescence lifetime, intensity, and running speed simultaneously as mice voluntarily ran or rested on a treadmill (Fig. 6A). For acute changes with one laser power and within one mouse, both intensity and lifetime of GRAB_ACh3.0_ showed an increase from resting to running, indicating that both properties capture transient ACh changes effectively (Fig. 6B). The increased intensity or lifetime from resting to running was not observed with the mutant sensor GRAB_ACh3.0mut_ (Fig. S6B-S6D).

**Figure 6.**
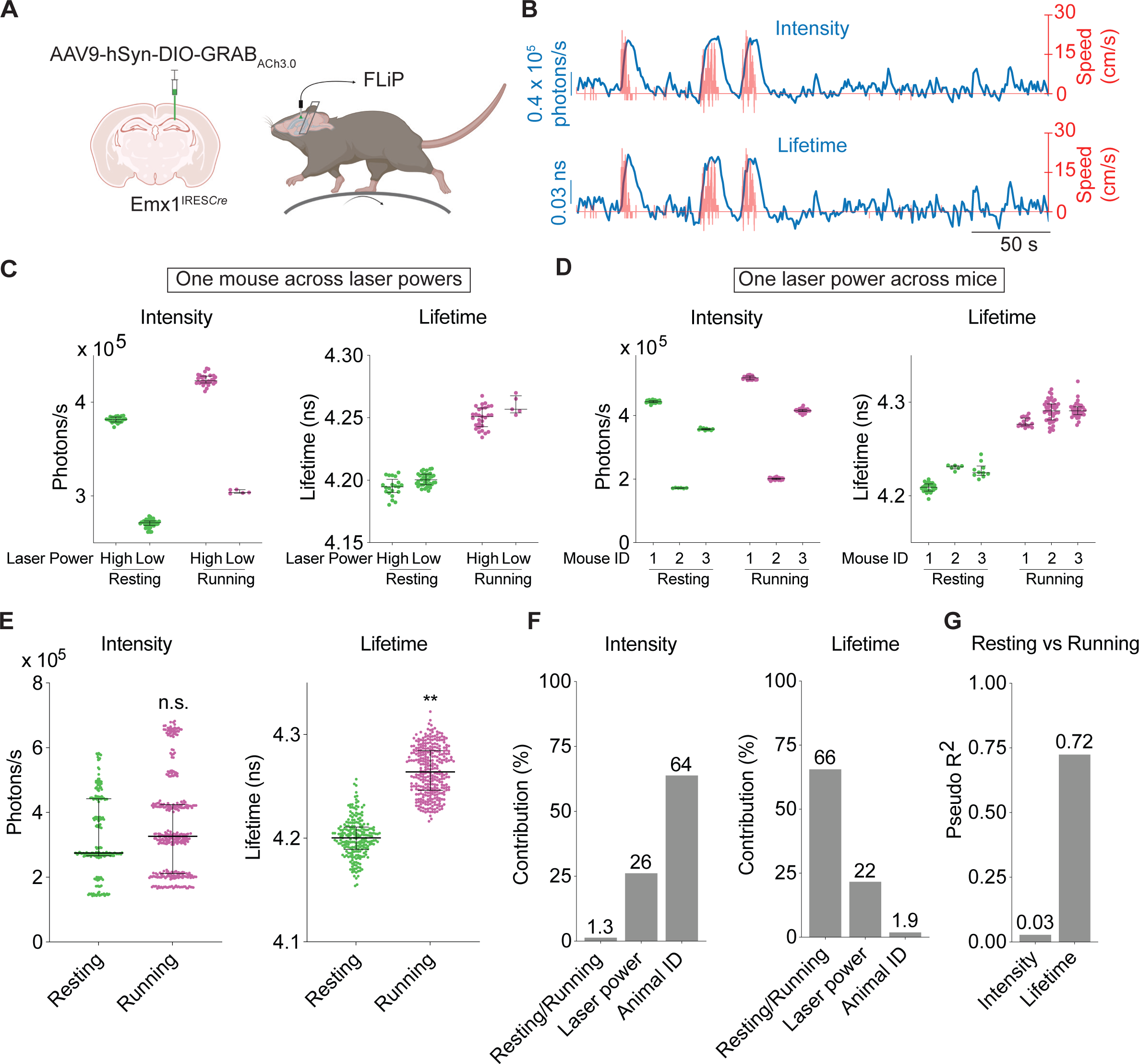
Fluorescence lifetime of GRAB_ACh3.0_ correlates with running vs resting states accurately despite varying laser powers and varying sensor expression levels across mice in vivo. **(A)** Schematic showing the experimental setup. AAV carrying Cre-dependent GRAB_ACh3.0_ was delivered to CA1 cells in the hippocampus of Emx1^IRES*cre*^ mice. FLiP was performed as head-fixed mice ran or rested on a treadmill. **(B)** Example traces showing intensity (top, blue) or fluorescence lifetime (bottom, blue) measurements from FLiP, and running speed (red) of GRAB_ACh3.0_-expressing mice on a treadmill. **(C)** Distribution of intensity and fluorescence lifetime of GRAB_ACh3.0_ in resting or running states from the same mouse but under different laser powers. **(D)** Distribution of intensity and fluorescence lifetime of GRAB_ACh3.0_ in resting or running states under the same laser power but from different mice. **(E)** Distribution of intensity and fluorescence lifetime of GRAB_ACh3.0_ in running or resting states, pooled from all mice across different laser powers (12 recordings from 6 mice under 3 different laser powers). Nested t test, **p < 0.01; n.s. not significant. **(F)** Results from stepwise-GLM analysis showing the contribution to the total variation of intensity or fluorescence lifetime of GRAB_ACh3.0_ from behavior states, laser power, and animal identities. Contribution is calculated from adjusted incremental R^2^. **(G)** Results from logistic regression analysis showing the power of explaining running or resting states with either intensity or fluorescence lifetime of GRAB_ACh3.0_, regardless of imaging laser powers or animal identities. Data are represented as median with interquartile range. See also Figure S6.

To test how well absolute values of lifetime or intensity correlates with ACh concentrations without information of transient changes, we asked how accurately we can explain running versus resting states across varying laser powers and across individual mice. These conditions mimic realistic scenarios because fluctuating laser power can arise from an unstable laser source or movement artifacts, and comparison across mice is essential when control versus disease models are compared.

Across varying laser powers, intensity showed large variation within the same resting or running state, whereas fluorescence lifetime remained remarkably stable (Fig. 6C). Similarly, with one laser power across different mice, intensity varied greatly within the same behavior state, likely due to different sensor expression level across mice. In contrast, lifetime remained stable within each running and resting state (Fig. 6D). When data from different imaging conditions and mice were combined, fluorescence intensity was not statistically different between running and resting (Fig. 6E; n = 226 resting epochs and 322 running epochs from 6 mice, p = 0.37), indicating that the absolute values of intensity could not be used to distinguish ACh levels between mice and between imaging conditions. Remarkably, despite these differing conditions, lifetime showed significant increase from resting to running (Fig 6E; p < 0.0001). These results indicate that in contrast to intensity, lifetime is stable across imaging power and across mice, and can distinguish ACh-associated behavior states across these conditions.

To quantitate the power of fluorescence lifetime, we performed two statistical tests. First, we asked how much of the variance of lifetime and intensity could be explained by running versus resting states, laser power, and animal identity. For fluorescence intensity, most of the variance was explained by animal identity (64%), followed by laser power fluctuation (26%), with minimal variance explained by behavior state (1.3%) (Fig. 6F, calculated from adjusted incremental R^2^ of stepwise generalized linear model (stepwise-GLM)). In contrast, most of the variance in lifetime was explained by behavior state (66%), with small contributions from laser power (22%) and animal identity (1.9%) (Fig. 6F, adjusted incremental R^2^ of stepwise-GLM). Secondly, we performed logistical regression to ask how much we could explain running versus resting state solely based on lifetime or intensity. Lifetime showed much better explanatory power than sintensity (Fig. 6G; pseudo R^2^ = 0.72 for lifetime and 0.03 for intensity). These results indicate that fluorescence lifetime, but not intensity, correlates with neuromodulator-associated behavior states despite fluctuating laser powers and expression level changes across animals.

Together, although both intensity and lifetime of GRAB_ACh3.0_ capture acute neuromodulator changes effectively, lifetime excels when experiments call for comparison of neuromodulator levels across fluctuating laser powers and across animals.

### Fluorescence lifetime correlates with ACh-associated sleep-wake states more accurately than intensity across chronic time scales

In vivo, the expression levels of a fluorescent sensor vary both across animals and across chronic time scales. We thus investigated whether fluorescence lifetime can accurately track ACh levels over many weeks, even as sensor expression levels change. We used sleep-wake cycles of mice as our proof-of-principle experiment because hippocampal ACh is higher during active wake (AW) and REM sleep, and low during quiet wake (QW) and NREM sleep^56, 64–69^. To evaluate the power of lifetime and intensity in explaining ACh-associated sleep and wake stages, we measured lifetime and intensity of the ACh sensor with FLiP in freely behaving mice, while simultaneously performing electroencephalogram (EEG), electromyography (EMG), and video recordings to determine sleep-wake stages (Fig. 7A).

**Figure 7.**
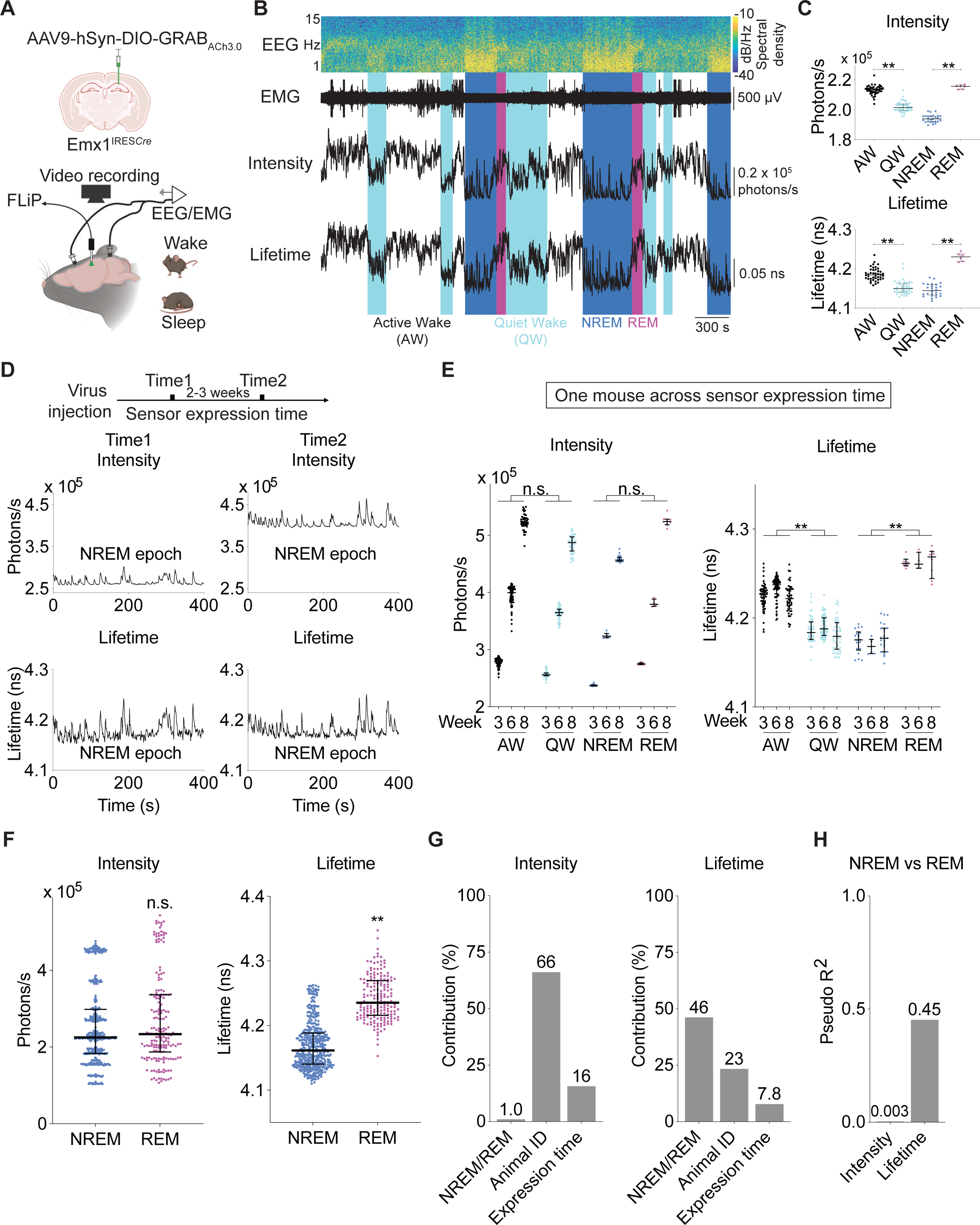
Fluorescence lifetime of GRAB_ACh3.0_ correlates with sleep-wake stages accurately despite variation in sensor expression levels across weeks and across animals. **(A)** Schematic showing the experimental setup. AAV carrying Cre-dependent GRAB_ACh3.0_ was delivered to CA1 cells in the hippocampus of Emx1^IRES*cre*^ mice. FLiP, EEG, EMG, and video recordings were performed across sleep-wake cycles over 9 hours (9 pm to 6 am) in freely moving mice. **(B)** Example of spectrogram of EEG recording, EMG trace, the corresponding scored sleep-wake states, along with intensity and fluorescence lifetime traces from a mouse within 1 hour. Note increases in GRAB_ACh3.0_ intensity and lifetime during REM and active wake. **(C)** Distribution of intensity and fluorescence lifetime of GRAB_ACh3.0_ in different sleep-wake states from a 9-hour FLiP recording of one mouse. Kruskal-Wallis test with Dunn’s multiple comparison, **p < 0.01. **(D)** Representative traces of intensity and fluorescence lifetime of GRAB_ACh3.0_ during NREM at two time points after virus injection. Note that fluorescence lifetime measurement was stable over time whereas intensity showed a large increase over time. **(E)** Summaries of intensity and fluorescence lifetime of GRAB_ACh3.0_ in different sleep-wake stages in one mouse across sensor expression time. Nested t test, **p < 0.01, n.s. not significant. **(F)** Distribution of intensity and fluorescence lifetime of GRAB_ACh3.0_ across NREM and REM sleep states, pooled from all mice across different sensor expression time (18 recordings from 6 mice at 3 different sensor expression time). Nested t test, **p < 0.01; n.s. not significant. **(G)** Results from stepwise-GLM analysis showing the contribution to the total variation of intensity or fluorescence lifetime of GRAB_ACh3.0_ from behavior states (NREM vs REM), sensor expression time, or animal identities. Contribution is calculated from adjusted incremental R^2^. **(H)** Results from logistic regression analysis showing the power of explaining NREM vs REM states with either intensity or fluorescence lifetime of GRAB_ACh3.0_, regardless of sensor expression time or animal identities. Data are represented as median with interquartile range. See also Figure S7.

We first asked whether lifetime, like intensity, reports acute changes of ACh as mice transition between different sleep-wake stages. For a given mouse recorded within a single day, both fluorescence lifetime and intensity of GRAB_ACh3.0_ increased from QW to AW, and from NREM to REM sleep (Fig. 7B-7C; n = 42, 42, 26, 6 epochs for AW, QW, NREM, and REM respectively; adjusted p < 0.0001 for AW vs QW and NREM vs REM of both intensity and lifetime). These results indicate that fluorescence lifetime, like intensity^56^, can detect acute ACh changes across sleep/wake stages.

Controlling for the specificity of the response, we performed the same experiment with the mutant ACh sensor GRAB_ACh3.0mut_ that does not bind to ACh (Fig. S7). Unexpectedly, GRAB_ACh3.0mut_ showed an acute decrease in fluorescence intensity as mice transitioned from NREM to REM sleep (Fig S7A-S7B; n = 42, 22, 50, 14 epochs for AW, QW, NREM, and REM respectively; adjusted p = 0.25 for AW vs QW and 0.0002 for NREM vs REM). Fluorescence lifetime did not show significant change between AW and QW, or between NREM and REM (Fig. S7B; adjusted p = 0.46 for AW vs QW and 0.51 for NREM vs REM). Because mutant ACh sensor responds to other environmental factors and not ACh, these data emphasize the importance of mutant sensor controls in the use of neuromodulator sensors.

To test the consistency of fluorescence lifetime as sensor expression level varies across long periods of time, after viral delivery of GRAB_ACh3.0_, we measured lifetime and intensity at three different time points that were weeks apart. As expected, fluorescence intensity showed drastic changes over time (Fig. 7D-7E). When results were pooled across sensor expression time, intensity values were not significantly different between different behavior states (Fig. 7E; n = 169, 152, 48, 18 total epochs for AW, QW, NREM, and REM respectively; p = 0.77 for AW vs QW, and 0.61 for NREM vs REM). In contrast, fluorescence lifetime remained stable for a given behavioral state, even as sensor expression changed over time (Fig. 7D-7E). Lifetime values were significantly different between behavior states despite sensor expression variation (Fig. 7E; p = 0.0007 for AW vs QW, and < 0.0001 for NREM vs REM). Therefore, these results indicate that fluorescence lifetime, unlike intensity, is stable as sensor expression changes over weeks, and is strongly correlated with ACh-associated behavior states.

To ask whether lifetime correlates with ACh-associated NREM/REM states despite varying sensor expression levels across chronic time scales and across mice, we combined results from different sensor expression time and mice. Lifetime, unlike intensity, was still significantly different between NREM and REM sleep states (Fig. 7F; n = 444 NREM epochs and 183 REM epochs from 6 mice; p = 0.72 for intensity and 0.0006 for lifetime).

To quantitate the contributions to variation of lifetime and intensity by different factors, we calculated adjusted incremental R^2^ from stepwise-GLM. The variation of fluorescence intensity was largely explained by animal identity (66%), followed by sensor expression time (16%), with minimal contribution from behavior states (1.0%) (Fig. 7G). In contrast, lifetime variation was largely explained by NREM versus REM states (46%), with much less contribution from animal identity (23%) and sensor expression time (7.8%; Fig. 7G).

Conversely, we tested the extent to which lifetime or intensity could distinguish ACh-associated sleep stages. Lifetime showed much higher explanatory power for NREM versus REM states than intensity, despite changing expression level and across different animals (Fig. 7H; pseudo R^2^ = 0.003 for intensity and 0.45 for lifetime). Therefore, fluorescence lifetime is a better correlate of behavior state than intensity, when data from multiple animals and across weeks need to be considered.

Taken together, these results indicate that in vivo, fluorescence lifetime, like intensity, captures acute changes in neuromodulator levels within one animal. Importantly, fluorescence lifetime, and not intensity, correlates with neuromodulator levels and has much greater explanatory power than intensity when experiments call for comparison between animals and across long periods of time.

## DISCUSSION

In summary, we discovered fluorescence lifetime responses for multiple neuromodulator sensors. Fluorescence intensity enables measurement of acute changes of neuromodulator levels at high temporal resolution. However, due to its sensitivity to laser power fluctuation and sensor expression levels, it is not suitable for making comparisons across days and across animals. Fluorescence lifetime measurement can overcome these limitations. Like fluorescence intensity, we found that fluorescence lifetime can detect transient neuromodulator changes and is dose sensitive. In contrast to fluorescence intensity, fluorescence lifetime is consistent and shows little variability with varying laser powers, with repeated measurements within a cell, and with different sensor expression levels between cells. In vivo, fluorescence lifetime, unlike intensity, still correlates with neuromodulator levels even as sensor expression level changes across days and across animals. Thus, fluorescence lifetime measurement of neuromodulator sensors opens doors to study neuromodulator dynamics at high spatial and temporal resolution across animals, brain regions, and chronic time scale (Fig. 8).

**Figure 8.**
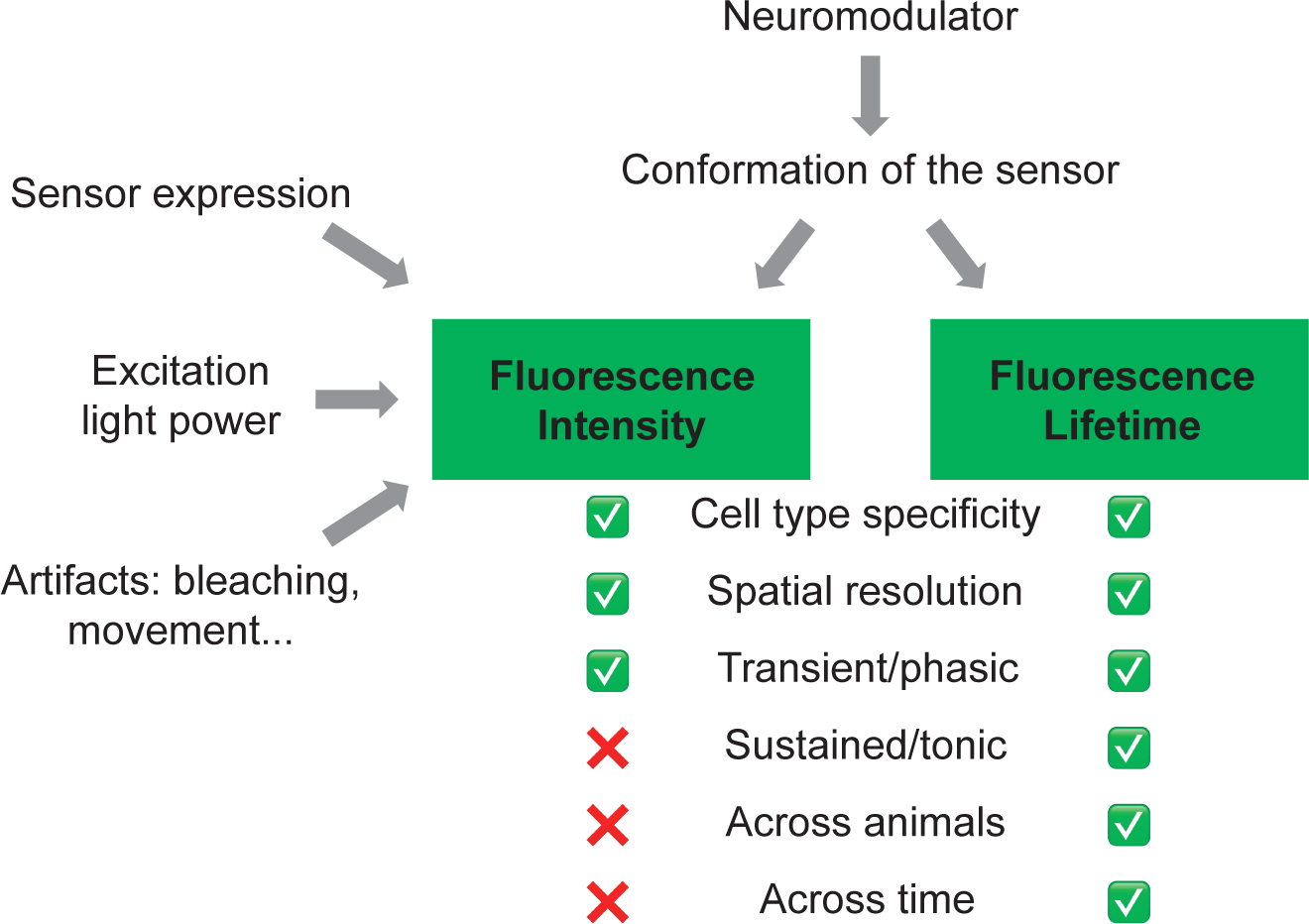
Comparison of intensity and lifetime measurement of fluorescent neuromodulator sensors. Fluorescence lifetime reflects conformation change of the sensor, whereas intensity is also influenced by sensor expression level, excitation light power, and other artifacts such as bleaching and movement. As a result, although fluorescence intensity excels in having cell type specificity, high spatial resolution, and high temporal resolution to detect transient/phasic changes of neuromodulators, it cannot be used to compare sustained/tonic changes of neuromodulators, compare neuromodulator levels across animals or chronic time scale. Fluorescence lifetime, in contrast, excels in all these categories.

### Advantages of using fluorescence lifetime to measure neuromodulator concentrations

When should we use lifetime over intensity measurement? Based on our results (Fig. 6 and 7), both lifetime and intensity can report acute neuromodulator changes. Fluorescence lifetime excels over intensity because lifetime measurement is independent of sensor expression^38, 40–43^. Due to this property, we demonstrate four major advantages of lifetime measurement in our proof-of-principle experiments. First, it is a robust correlate of neuromodulator concentration despite changing sensor expression levels across individual animals (Fig. 6 and 7). Second, lifetime is stable despite fluctuating excitation light power (Fig. 4 and 6). Third, lifetime correlates with neuromodulator concentration with high accuracy despite large variation of sensor expression levels over chronic time scale of weeks (Fig. 7). Finally, as demonstrated in our mutant sensor data, fluorescence lifetime is not as sensitive as intensity to neuromodulator-independent change associated with NREM to REM transitions (Fig. S7). This REM-associated intensity decrease calls for careful interpretation of data to distinguish neuromodulator change from brain state-associated change in intensity measurement such as hemodynamic change. In summary, fluorescence lifetime excels over intensity when one needs to compare changes across individual animals, across fluctuating excitation light power, and across chronic time scale.

### Opportunities for new biology

The discovery and demonstration of the power of fluorescence lifetime-based sensors provide new opportunities for biological discovery (Fig. 8). As demonstrated in our proof-of-principle experiments with sleep-wake stages and running-resting states (Fig. 6 and 7), lifetime value is a much better correlate of neuromodulator concentration than intensity, enabling comparison of neuromodulator levels at high temporal resolution across changing light levels, between individual animals, and at different time points that are weeks or months apart. In addition, because lifetime is robust over varying sensor expression levels, it enables investigation of how neuromodulator levels differ between brain regions, between young and old animals during aging, and between control and disease models of neurological and psychiatric disorders. Furthermore, a fundamental yet unanswered question in neuromodulator research is whether phasic/transient or tonic/sustained change of neuromodulator release is the predominant driver between control and disease conditions, and in response to therapeutic drug treatment. Lifetime offers the opportunity to disambiguate transient and sustained change, a feat that neither fluorescence intensity measurement nor microdialysis can accomplish alone. Thus, lifetime measurement of neuromodulators holds exciting potential for studying normal physiology, disease processes, and drug effects.

### Opportunities for new sensor design

Current neuromodulator sensors have not been optimized for lifetime measurement because they have generally been selected for low intensity during baseline conditions, making lifetime measurement challenging. To optimize for lifetime response, sensors need to be screened for 1) increased brightness to make measurement of fluorescence lifetime reliable, 2) larger dynamic range between different neuromodulator concentrations, and 3) minimal variation in lifetime readout with the same neuromodulator concentration between cells and between animals. Despite the lack of optimization for fluorescence lifetime measurement, lifetime of GRAB_ACh3.0_ shows high signal-to-noise ratio and clear separation of behavior states in vivo (Figs. 6 and 7). Thus, our discovery of lifetime change by single fluorophore-based GPCR sensors provides the foundation and inspiration for developing more lifetime-based neuromodulator sensors. Given the demonstrated power of fluorescence lifetime for comparison across animals, between disease models, and across chronic time periods, all sensor developers should look at fluorescence lifetime, in addition to intensity, as a criterion for sensor screening and optimization in the future.

## AUTHOR CONTRIBUTIONS

Conceptualization: P.M. and Y.C.; Methodology: P.M., P.C., E.T. and Y.C.; Formal Analysis: P.M., P.C., S.A., and A.O.; Investigation: P.M., A.O., and Y.C.; Writing: P.M. and Y.C.; Visualization: P.M., A.O. and Y.C.; Supervision: Y.C.; Funding Acquisition: Y.C.

## ACKNOWLEDGEMENTS

We thank Yulong Li and lab for sharing plasmids of neuromodulator sensors and for discussions. We thank Sophie Ma for validation of sleep scoring results. We thank Adam Kepecs, Meaghan Creed, and the labs of Yao Chen, Tim Holy, and Daniel Kerschensteiner for helpful feedback on the project. We thank Martha Bagnall, Yanchao (Miko) Dai, Kerry Grens, Yulong Li, Aditi Maduskar, and Thomas Papouin for critical comments on the manuscript. Schematic illustrations from Figure 1A, 3A, 6A, 7A, and S6B were created with BioRender. Funding for this work was supported by the U.S. National Institute of Neurological Disorders and Stroke (R01 NS119821, to Y.C.), the Whitehall Foundation (2019-08-64, to Y.C.), a gift from the Howard Hughes Medical Institute (to Y.C.), and the McDonnell International Scholars Academy of Washington University in St. Louis (to P.M.).

## KEY RESOURCES TABLE

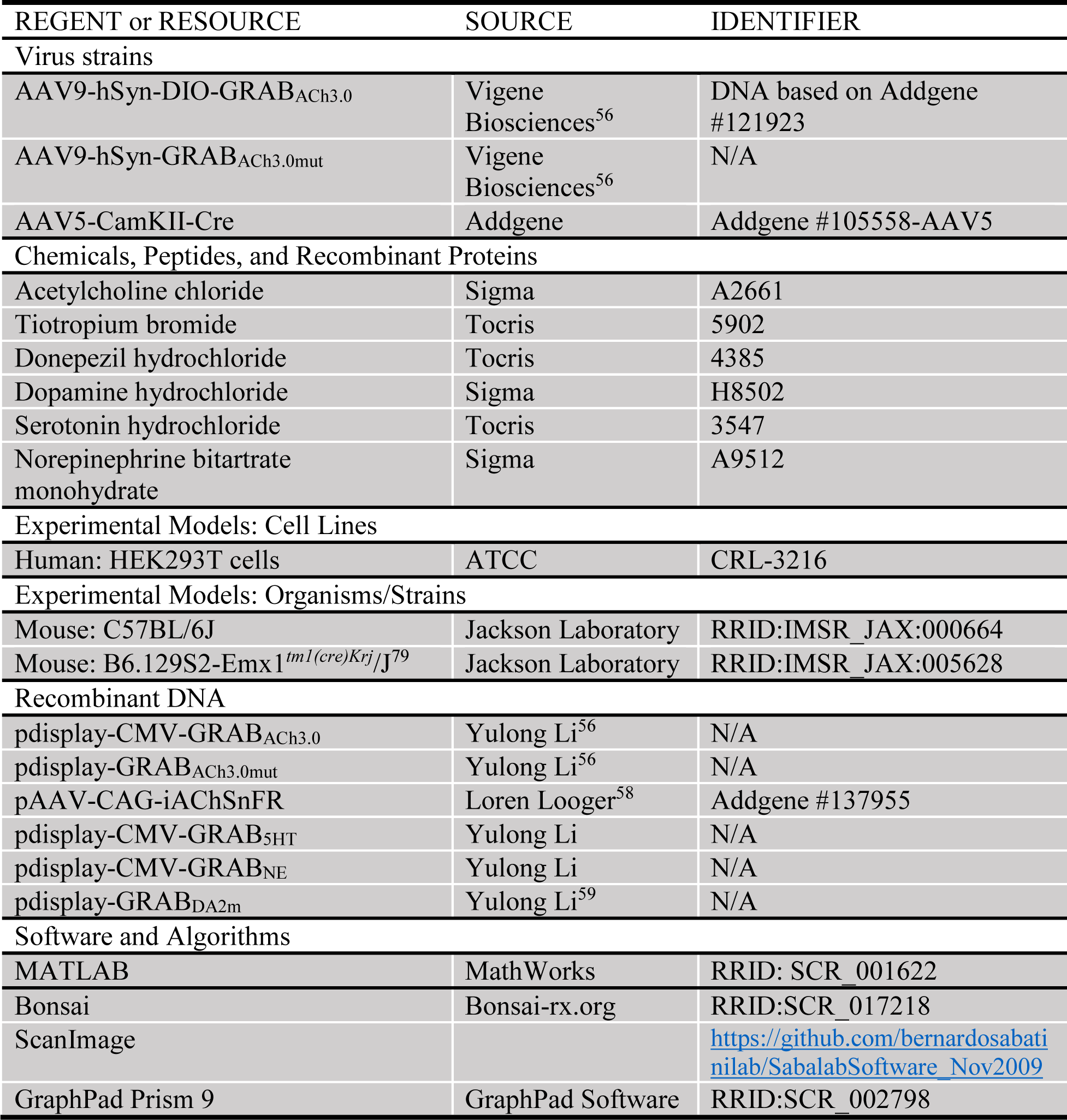

## STAR METHODS

### RESOURCE AVAILABILITY

#### Lead Contact

Further information and requests for resources and reagents should be directed to and will be fulfilled by the lead contact, Yao Chen.

#### Material Availability

This study did not generate new unique reagents.

### EXPERIMENTAL MODEL AND SUBJECT DETAILS

#### Human Embryonic Kidney (HEK) 293T Cells

HEK 293T cells were cultured in Dulbecco’s Modified Eagle Medium (DMEM) with 10% Fetal Bovine Serum (FBS) (Millipore Sigma), GlutaMAX (Invitrogen), and penicillin /streptavidin (50 U/m, Corning) at 37°C in 5% CO_2_. All cells were female. The cell line has not been authenticated. They were plated on coverslips in 24-well plates and transfected with plasmids (0.4-0.8 μg/well) using lipofectamine 2000 (Invitrogen). Two days after transfection, the cells were imaged with perfusion of artificial cerebrospinal fluid (ACSF, concentrations in mM: 127 NaCl, 25 Na_2_CO_3_, 1.25 NaH_2_PO_4_.H2O, 2.5 KCl, 1 MgCl_2_, 2 CaCl_2_, and 25 glucose).

#### Animals

All procedures for rodent husbandry and surgery were performed following protocols approved by the Washington University Institutional Animal Care and Use Committee and in accordance with National Institutes of Health guidelines. For acute brain slices, adult wild-type C57BL/6J mice (Jax 000664) were used with injections of virus expressing Cre recombinase and Cre-dependent sensors. For behavioral studies, adult Emx1^IRES*Cre*^ (Jax 005628) or wild-type mice were injected with virus and implanted with fiber-optic cannula, EEG/EMG implants, and headplates.

### METHODS DETAILS

#### DNA Plasmids

The constructs pdisplay-CMV-GRAB_ACh3.0_^56^, pdisplay-CMV-GRAB_5HT_, pdisplay-CMV-GRAB_NE_, pdisplay-GRAB_ACh3.0mut_^56^, and pdisplay-GRAB_DA2m_^59^ were gifts from Dr. Yulong Li’s laboratory. pAAV-CAG-iAChSnFR (Addgene #137955) was from Dr. Loren Looger’s laboratory^58^.

#### Virus Production and Stereotaxic Injections

AAV9-hSyn-DIO-GRAB_ACh3.0_^56^ (DNA corresponding to Addgene #121923) and AAV9-hSyn-GRAB_ACh3.0mut_^56^ viruses were packaged at Vigene Biosciences. AAV5-CamKII-Cre was from James M. Wilson and packaged at Addgene (Addgene #105558-AAV5). For stereotaxic injection, dorsal hippocampus CA1 was targeted with coordinates of posterior 1.78 mm and lateral 1.58 mm relative to Bregma, and 1.36 mm from the pia. All injections were made at a rate of 100 nL/min through a UMP3 micro-syringe pump (World Precision Instruments) via glass pipette. For acute brain slice imaging, bilateral injections of 500 nL of AAV9-hSyn-DIO-GRAB_ACh3.0_ (3.1 x 10^12^ GC/mL) and AAV5-CamKII-Cre (3 x 10^12^ GC/mL) were made in wild-type mice. For FLiP experiments, 500 nL of AAV9-hSyn-DIO-GRAB_ACh3.0_ (3.9 x 10^12^ GC/mL) were injected into left hemispheres of Emx1^IRES*Cre*^ mice. For control experiments, 500 nL of AAV9-hSyn-GRAB_ACh3.0mut_ (3.1 x 10^12^ GC/mL) were injected into the left hemispheres of wild-type mice. Following virus injection, fiber-optic cannula, EEG/EMG implants, and headplates were placed.

#### Implantation of Optic Cannula, EEG/EMG Implants, and Headplate

After stereotaxic injection and withdrawal of the glass pipette, a fiber-optic cannula (Doric Lenses, MFC_200/245-0.37_2.5mm_MF1.25_FLT) was inserted into the same injection site, at 0.05 mm above the viral injection site. The fiber was stabilized to the skull with glue. To implant the EEG and EMG implants, four stainless steel screws were inserted into the skull, with two above the cerebellum, one above the right hippocampus, and one above the right frontal cortex. The screws were wired to an EEG/EMG head-mount (Pinnacle, 8402). Two EMG electrodes from the head-mount were inserted into the neck muscle of the mice. A headplate was placed directly onto the skull. All the implants were secured to the skull with dental cement. An additional layer of dental cement with black paint was applied for light-proofing. All experiments were carried out at least 2 weeks after the surgery.

#### Acute Brain Slice Preparation

Mice were anesthetized with isoflurane followed by intracardial perfusion with cold N-methyl-d-glucamine (NMDG)-based cutting solution (concentrations in mM: 92 NMDG, 2.5 KCl, 1.25 NaH_2_PO_4_, 30 NaHCO_3_, 20 HEPES, 25 glucose, 10 MgSO_4_, 0.5 CaCl_2_, 5 sodium ascorbate, 2 thiourea, and 3 sodium pyruvate)^80^. Their brains were rapidly dissected out. 300 μm-thick coronal sections were obtained with a vibratome (Leica Instruments, VT1200S) in cold NMDG-based cutting solution. After sectioning, slices were transferred to NMDG-based solution and incubated at 34℃ for 12 minutes, and then kept in HEPES-based holding solution (concentrations in mM: 92 NaCl, 2.5 KCl, 1.25 NaH_2_PO_4_, 30 NaHCO_3_, 20 HEPES, 2 thiourea, 5 sodium ascorbate, 3 sodium pyruvate, 2 CaCl_2_, 2 MgSO_4_, and 25 glucose) at room temperature with 5% CO_2_ and 95% O_2_. Slices were then transferred to a microscope chamber and ACSF was perfused at a flow rate of 2-4 mL/min for imaging.

#### Two-Photon Fluorescence Lifetime Imaging (2pFLIM) and Image Analysis

Two photon imaging was achieved by a custom-built microscope with a mode-locked laser source (Spectra-Physics, Insight X3 operating at 80 MHz). Photons were collected with fast photomultiplier tubes (PMTs, Hamamatsu, H10770PB-40). A 60X objective (Olympus, NA 1.1) was used. Image acquisition was performed with the custom-written software ScanImage in MATLAB 2012b^81^.

FLIM was performed as described previously^48, 49^. For all the GFP-based neuromodulator sensors, 920 nm was used as the excitation wavelength. Emission light was collected through a dichroic mirror (FF580-FDi01-25X36, Semrock) and a band-pass filter (FF03-525/50-25, Semrock). 128x128 pixel images were collected by frame scan at 4 Hz. The FLIM board SPC-150 (Becker and Hickl GmbH) was used, and time-domain single photon counting was performed in 256 time channels. For FLIM data analysis, only healthy cells (judged by gradient contrast images) with membrane expression pattern were selected. Cells with round shape, sensor expression aggregates, or cell-filling expression patterns were excluded. The membrane of individual cells was selected as region of interest (ROI). To minimize the effect of movement artifact on intensity measurement, pixels with photon counts below 5 was omitted and then the top 66% brightest pixels were selected as effective pixels. Photons from effective pixels of a given ROI were pooled. The average photon count per pixel was used for intensity measurement. The average lifetime of all the photons in this ROI was calculated as follows:

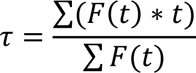

in which F(t) is the photon count from the fluorescence lifetime histogram at time bin t, and t is the lifetime measurement corresponding to the time bin. We performed the calculation from 0.0489 ns to 11.5 ns in the lifetime histogram. Due to change of cable length in FLIM or FLiP set-up, the empirical lifetime across different experiments showed different absolute values. The cable length was kept consistent within one set of experiments.

Change of fluorescence lifetime at baseline was quantitated as lifetime measurement averaged over the first 5 data points of baseline subtracted from lifetime measurement averaged over the last 5 data points of baseline. Change of lifetime due to treatment was calculated as the average lifetime of the last 5 data points of baseline subtracted from that of the last 5 data points of treatment period. Cells with unstable baseline (coefficient of variation for baseline lifetime larger than 0.8%) were excluded. Similar calculations were performed for intensity change, with change of intensity divided by the average intensity of the first 5 data points of baseline as ΔF/F_0_.

For puffing experiments, maximum of either lifetime or intensity during baseline or puffing period was used to calculate the response. For dose-dependent response experiments, the response of each concentration of ACh treatment was expressed as the percentage of the peak responses.

#### Fluorescence Lifetime Photometry (FLiP) and Analysis

A FLiP setup was custom built and used similar to that previously described^78^. In brief, a pulsed 473nm laser (Becker and Hickl, BDS-473-SM-FBE operating at 50 MHz) was used as the excitation light source. An optical fiber patch cord (Doric Lenses, MFP_200/220/900-0.37_1.5m_FCM-MF1.25_LAF) was used to direct the excitation laser beam to the optical fiber implanted in the mouse brain. A dichroic mirror (Thorlabs, DMLP505R) and band-pass filter (Semrock, FF01-525/39-25) were used to select the green emission light from the blue excitation light. Emission light was detected with a fast photomultiplier tube (PMT, Hamamatsu, H10770PA-40), and a time-correlated single-photon counting (TCSPC SPC-150, Becker and Hickl GmbH) board was used to measure fluorescence lifetime binned into 256 time channels. The data were collected by customized software in MATLAB 2012b at 1 Hz. Excitation light power was adjusted with a neutral density filter, so the photon arrival rate was between 1 x 10^5^/s and 8 x 10^5^/s. The lower limit was chosen for accurate estimation of lifetime, and the upper limit chosen based on the dead time of the TCSPC driver board. The typical excitation power needed to generate the appropriate rate of photons for TCSPC was 0.01–0.18 μW (measured at the output end of the patch cord). Location of viral injection and fiber implants examined by histology after experiments. Only mice with tip of the fiber above hippocampus CA1 were used in the behavior analysis. For data analysis, we calculated average lifetime from 2.148 ns to 18.555 ns in the lifetime histogram.

#### Running and Resting Recording and Analysis

Mice with optic fiber implant and headplate were head-fixed on a treadmill and recorded in the dark. An incremental rotary encoder (SparkFun, COM-11102) was used to record the speed of the voluntary running. Rotary signals were collected at 25Hz via an Arduino Due board (Arduino, A000062). The signals were sent to Bonsai (https://bonsai-rx.org/) via serial port communication and timestamped in Bonsai. Videos were simultaneously recorded at 25 frames per second (fps) in Bonsai. FLiP data were collected at 1 Hz.

Raw data of running speed were binned to 4 Hz for analysis. Running epochs were defined by the following criteria: 1) continuous forward or backward movement above a speed of 1cm/s; 2) no more than 3 consecutive sub-threshold data points; 3) preceded by at least 10 seconds of sub-threshold resting; and 4) at least 3 seconds in duration. For ACh sensor fluorescence analysis during running, in order to account for sensor kinetics, 0.5 second at the beginning of each running epoch was excluded and 2 seconds were extended from the end of the running epochs for analysis Each resting epoch was specified as continuous below-threshold speed that lasts for more than 150 seconds. To account for sensor kinetics and ACh kinetics, the first and last 30 seconds of each resting epoch were excluded for analysis. If a trimmed resting epoch is longer than 90 seconds, it is split into 90 second epoch segments.

The median values of fluorescence intensity or fluorescence lifetime of ACh sensor for each running or resting segment were quantitated for subsequent analysis.

#### FLiP, EEG/EMG, and Video Recordings

Mice that underwent GRAB_ACh3.0_ virus injection, optical fiber implantation, and EEG/EMG implant were placed in a chamber with 12-hour/12-hour light-dark cycle (6am-6pm light). Recordings from 9pm to 6 am (dark phase) were collected and analyzed. An additional infrared light was used for video recording during the dark phase. Fluorescence lifetime and intensity data were collected at 1 Hz with our custom-built FLiP setup. EEG/EMG recording was performed at 400 Hz with a system from Pinnacle Technology using our ScanImage software. Video recording was performed at 25 fps in Bonsai. Video data were synchronized with FLiP and EEG/EMG data via a TTL (transistor-transistor logic) signal from Matlab to Arduino Due board (Arduino, A000062) to Bonsai to trigger the start of video recording.

#### Sleep Stage Scoring

Sleep stages were scored for every 4-second bin based on the EEG, EMG, and motion detection from the video using a custom-written program in Python. Briefly, sleep scoring prediction was generated with a random forest model, followed by user correction. The following criteria were used to determine sleep/wake stages^56, 82^: 1) active wake: low variance in EEG, high variance in EMG, and high movement based on video; 2) quiet wakefulness: low variance in EEG, low variance in EMG, and low movement based on video; 3) NREM sleep: high variance in EEG with high delta power (0.5-4 Hz), low variance in EMG, and no movement based on video; 4) REM sleep: high theta (5-8 Hz) to delta power ratio based on EEG, low variance in EMG, and no movement based on video.

#### Pharmacology

Unless otherwise noted, all chemicals were applied via bath perfusion: they were either added to the perfusion reservoir or pre-made buffers with the specified chemicals were switched from one to another. Lifetime was allowed to stabilize before a new chemical was added. When there was no clear lifetime change, 10 minutes were recorded before the addition of another chemical or the end of the experiment. The final concentrations of chemicals are specified in parentheses: ACh chloride (0.001 μM to 100 μM), norepinephrine bitartrate monohydrate (NE, 10 μM) and dopamine hydrochloride (DA, 10 μM) were from Sigma; serotonin hydrochloride (5-HT, 100 μM), muscarinic acetylcholine receptor antagonist tiotropium bromide (Tio, 5 μM), and cholinesterase inhibitor donepezil hydrochloride (5 μM), were from Tocris.

For puffing experiments, a glass patch pipette was used to locally puff ACh (200 μM in ACSF) for 3 seconds onto a neuron in a brain slice through a Picospritzer (Parker, 052-0500-900) at 2 psi.

#### FLIM Simulation

The simulation was performed by customized MATLAB code. The null hypothesis is that with or without ACh binding, GRAB_ACh3.0_ has the same fluorescence lifetime and can be described by the same equation – thus, the apparent fluorescence lifetime change was solely due to altered proportion of autofluorescence contribution. The fluorescence of GRAB_ACh3.0_ was modelled by a double exponential decay.

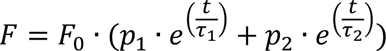

τ1, τ2, p1, and p2 were determined empirically by measuring the fluorescence decay of ACh 3.0 expressed in HEK cells at saturating concentration (100 μM) of ACh. A large population of photons (∼6 x 10^6^) with specific lifetimes was generated based on the double exponential decay and binned into 256 time channels over 12.5 ns (time interval between laser pulses for an 80 MHz laser). The photon population was then convolved by a pulse response function measured empirically. A small sample of photons was drawn with replacement from the large population, and the number of photons in the sample corresponded to the average of measured photons at either 0 or 100 µM of ACh respectively. Subsequently, we added to the photon sample photons due to afterpulse (0.32% of total photon count, with even distribution across lifetime) and autofluorescence. Photons due to autofluorescence were sampled from empirically determined autofluorescence distribution from untransfected HEK 293T cells. Simulation was repeated 500 times for each sample size corresponding to 0 or 100 µM of ACh. Empirical fluorescence lifetime was calculated for each simulated combination and compared to experimentally observed values.

### QUANTIFICATION AND STATISTICAL ANALYSIS

Detailed information of the quantification and statistics used are summarized in Figure Legends, Figures, and Results. Wilcoxon test (with Bonferroni correction when appropriate) was performed for paired data. Mann-Whitney test was performed for unpaired data. For analysis of variance, Friedman test was performed for matched data, and Kruskal-Wallis test was performed for unmatched data, followed by Dunn’s multiple comparison (one-way ANOVA), or Šídák’s multiple comparison (two-way ANOVA). Nested t test was performed when comparison was made with hierarchical data. All statistical analysis were performed in GraphPad Prism 9. Two-way ANOVA was used to determine the contribution to the total variance from two independent variables. When more than two independent variables were included, a stepwise-GLM model was performed in MATLAB. The independent variables were added in order of weights (largest first based on adjusted R^2^) and the subsequent improvement to overall adjusted R^2^ was calculated as the contribution to the variance for each independent variable. Logistic regression (LR) was used to identify the strength of the relationship of individual independent variables (intensity and lifetime) on states (resting/running; REM/NREM). LR was performed using Scikit-Learn in Python. McFadden’s pseudo R^2^ values were used to evaluate the performance of the model. Sample size n refers to biological replicates of number of cells, mice, or behavioral epochs.

### DATA AND SOFTWARE AVAILABILITY

The MATLAB programs for ScanImage for data acquisition and analysis are available at https://github.com/YaoChenLabWashU/2pFLIM_acquisition. The MATLAB codes for simulation are available at https://github.com/YaoChenLabWashU/Simulation. The Python codes for analysis of running vs resting states are available at https://github.com/YaoChenLabWashU/RVR_v2/. The Python codes for sleep staging are available at https://github.com/YaoChenLabWashU/neuroscience_sleep_scoring. Any additional information required to reanalyze the data reported in this paper is available from the lead contact upon request.

## SUPPLEMENTAL FIGURE LEGENDS

**Figure S1.**
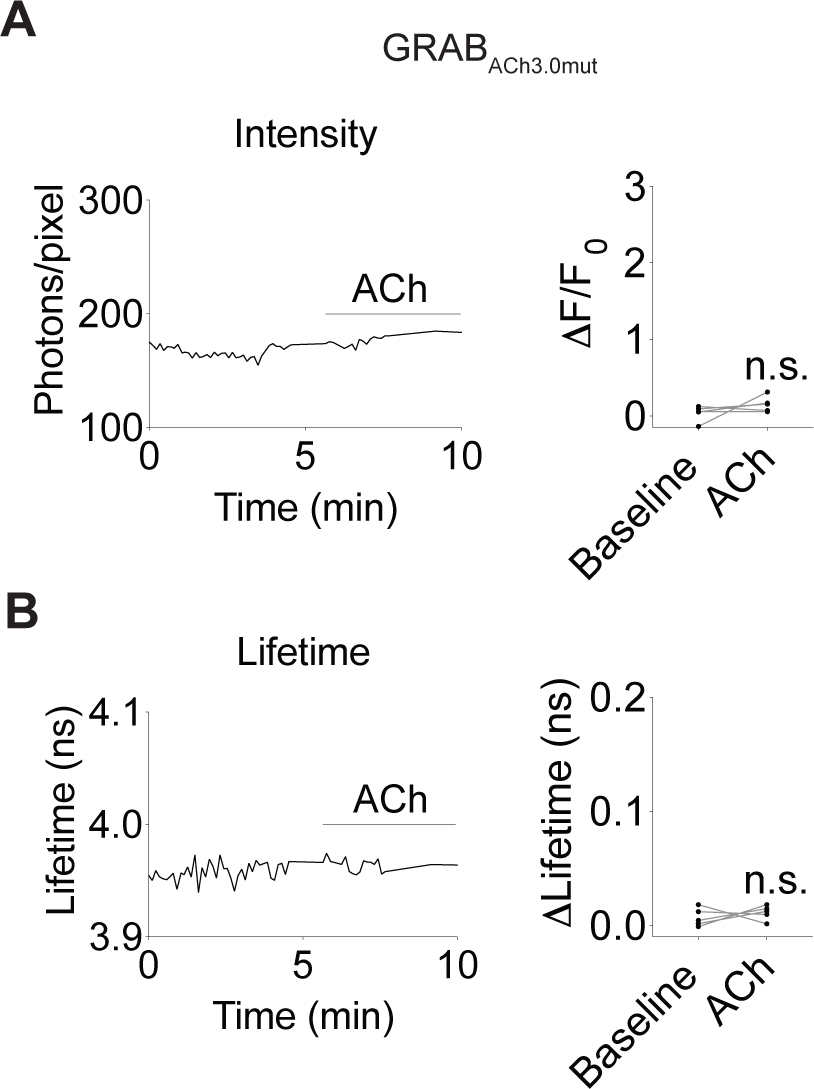
GRAB_ACh3.0mut_ sensor showed no fluorescence lifetime or intensity change to ACh application. **(A-B)** Traces (left) and summaries (right) of intensity (A) and fluorescence lifetime (B) response of GRAB_ACh3.0mut_ sensor to ACh application (100 µM). Wilcoxon test, n.s., not significant vs baseline.

**Figure S2.**
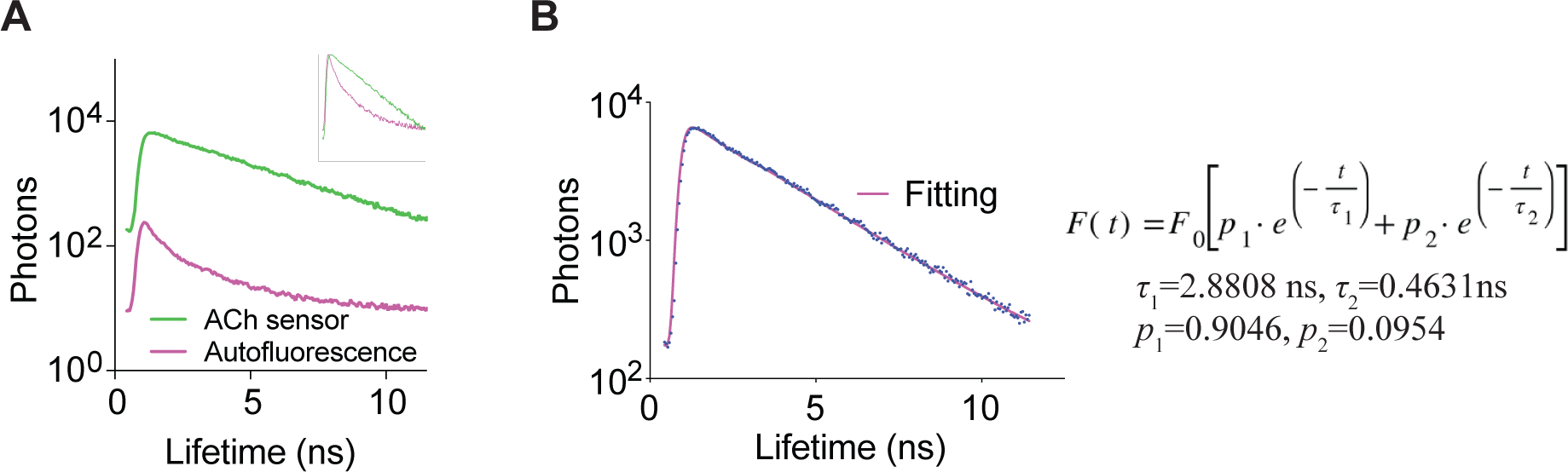
Fluorescence lifetime of sensor fluorescence and autofluorescence. **(A)** Measured histogram of fluorescence lifetime of HEK293T cells without sensor expression (autofluorescence), and with GRAB_ACh3.0_ expression (in the presence of 100 μM ACh). The inset shows histogram normalized to peak photon count, indicating that the fluorescence lifetime of autofluorescence is shorter than sensor fluorescence. **(B)** Measured histogram and corresponding double exponential curve fitting results of fluorescence lifetime of GRAB_ACh3.0_ in HEK293T cells in the presence of 100 μM ACh.

**Figure S3.**
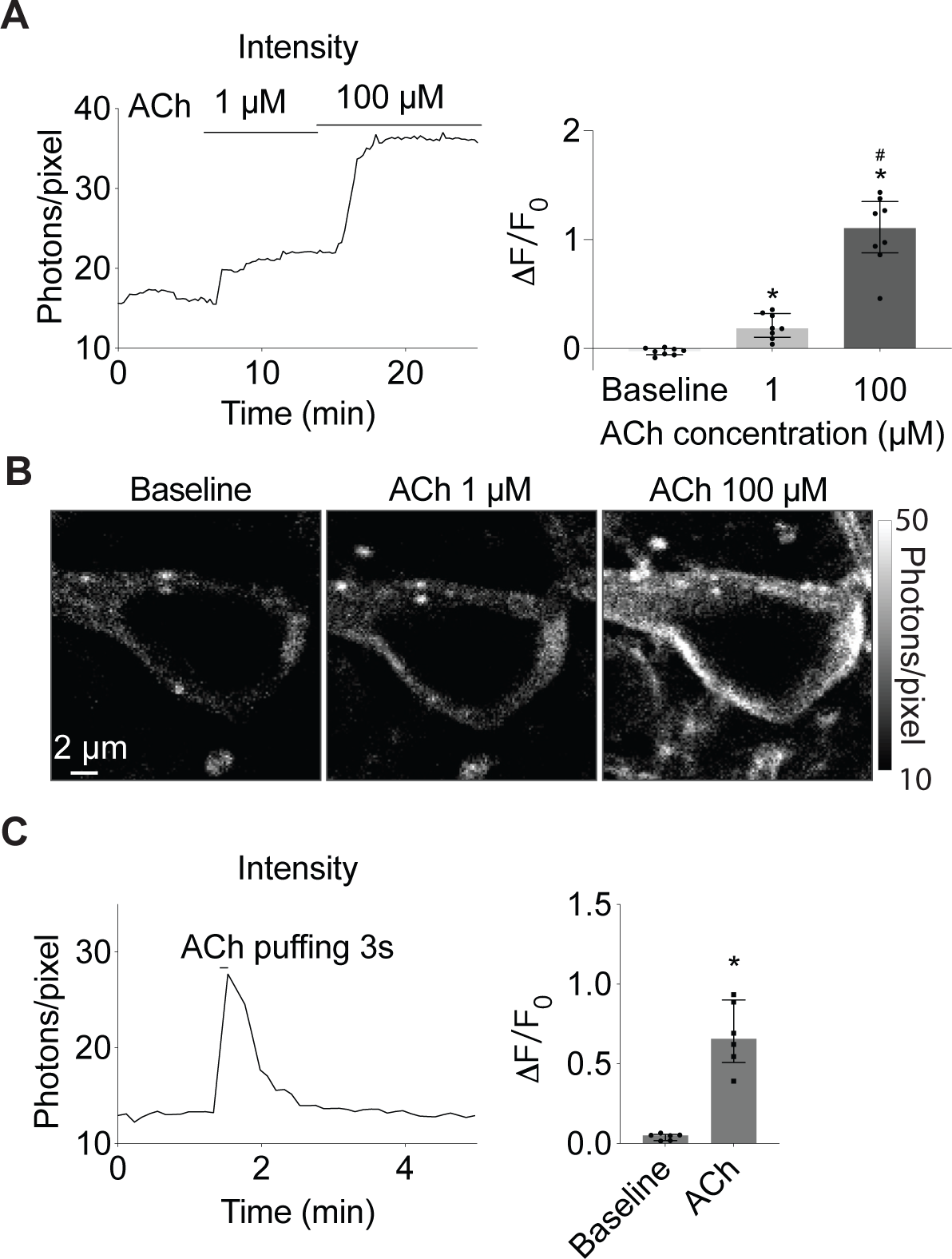
Intensity of GRAB_ACh3.0_ responds to transient ACh in brain tissue. **(A-B)** Example trace and summaries (A), as well as heatmap (B) showing intensity of hippocampal CA1 pyramidal neurons expressing GRAB_ACh3.0_ in response to ACh (1 µM and 100 µM, with 5 µM AChEi Donepezil). Wilcoxon test with Bonferroni correction, *p < 0.05 vs baseline, ^#^p < 0.05 vs 1 µM. **(C)** Example trace and summaries showing fluorescence intensity of GRAB_ACh3.0_ in CA1 pyramidal neurons in response to a 3-second puff of ACh (200 μM). Wilcoxon test, *p < 0.05 vs baseline. Data are represented as median with interquartile range.

**Figure S4.**
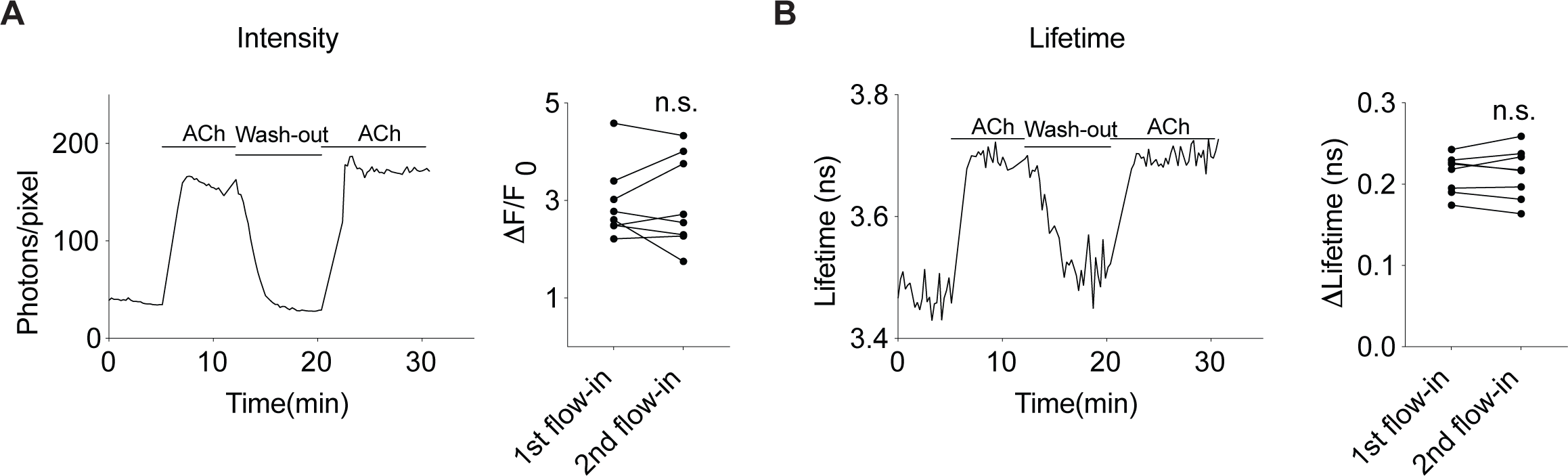
Fluorescence lifetime measurement of GRAB_ACh3.0_ is stable with repeated ACh application. **(A-B)** Trace and summaries of intensity (A) and fluorescence lifetime (B) measurement of GRAB_ACh3.0_ in HEK 293T cells in response to repeated flow-in of the same concentration of Ach (1 μM, with 5 μM of AChEi Donepezil). Wilcoxon test; n.s.: not significant, vs 1^st^ flow-in. Data are represented as median with interquartile range.

**Figure S5.**
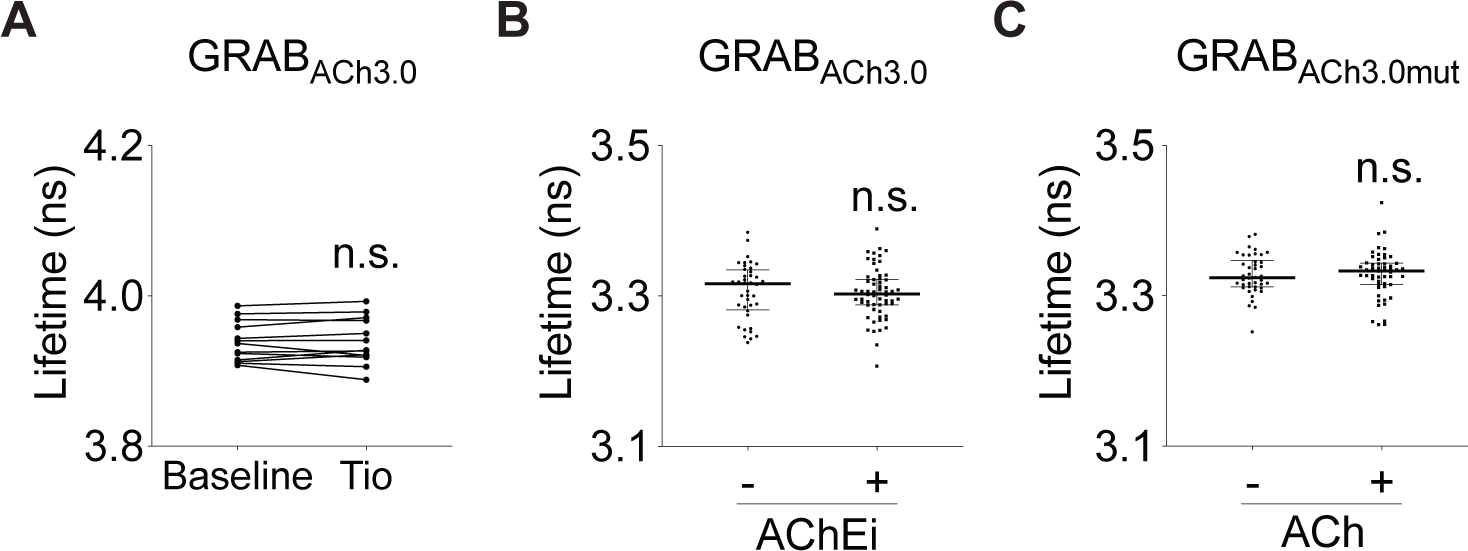
Distribution of fluorescence lifetime of GRAB_ACh3.0_ and GRAB_ACh3.0mut_. **(A)** Fluorescence lifetime of GRAB_ACh3.0_ in HEK 293T cells before and after application of mAChR antagonist Tiotropium (5 μM). Wilcoxon test, n.s., not significant vs baseline. **(B)** Distribution of fluorescence lifetime of GRAB_ACh3.0_ in HEK 293T cells with or without AChEi Donepezil (5 μM). Mann-Whitney test, n.s., not significant vs without AChEi. **(C)** Distribution of fluorescence lifetime of GRAB_ACh3.0mut_ sensor in HEK 293T cells with or without ACh (100 μM). Mann-Whitney test, n.s., not significant vs without ACh. Data are represented as median with interquartile range.

**Figure S6.**
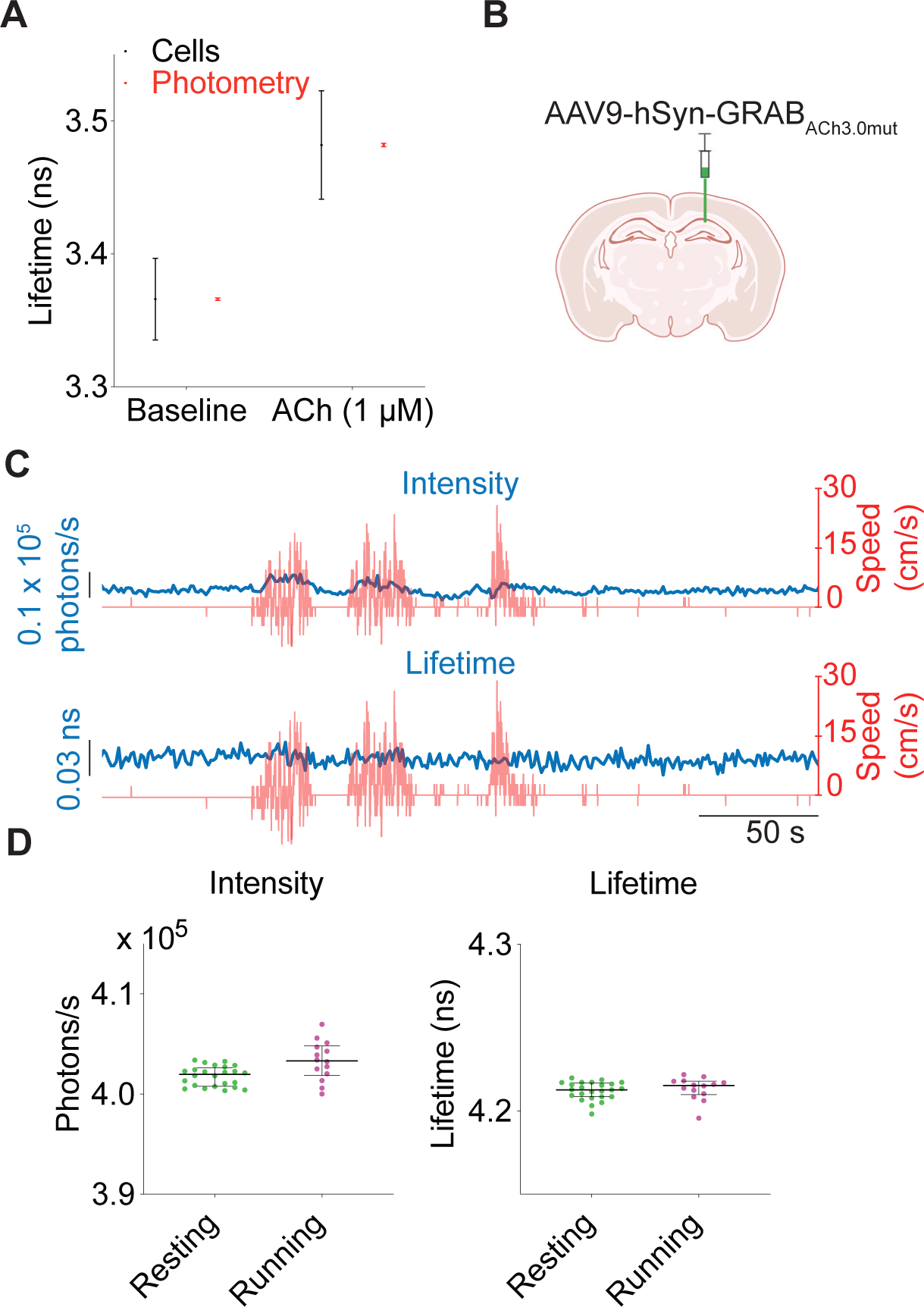
Intensity and fluorescence lifetime measurements of GRAB_ACh3.0mut_ sensor in the hippocampus during running/resting. **(A)** Schematic illustrating the reduction in standard deviation of data in bulk measurement with FLiP compared with cell-based imaging. Photometry data were modelled based on light collection from 1000 cells. Data are represented as mean with standard deviation. **(B)** Illustration showing expression of GRAB_ACh3.0mut_ in CA1 cells of hippocampus. AAV carrying GRAB_ACh3.0mut_ driven by neuronal specific hSyn promoter was delivered to CA1 cells in the hippocampus of wild-type mice. **(C)** Example traces showing intensity (top, blue) or fluorescence lifetime (bottom, blue) measurements from FLiP, and running speed (red) of GRAB_ACh3.0mut_-expressing mice on a treadmill. **(D)** Distribution of intensity and fluorescence lifetime of GRAB_ACh3.0mut_ sensor in resting or running states of one mouse. Data are represented as median with interquartile range.

**Figure S7.**
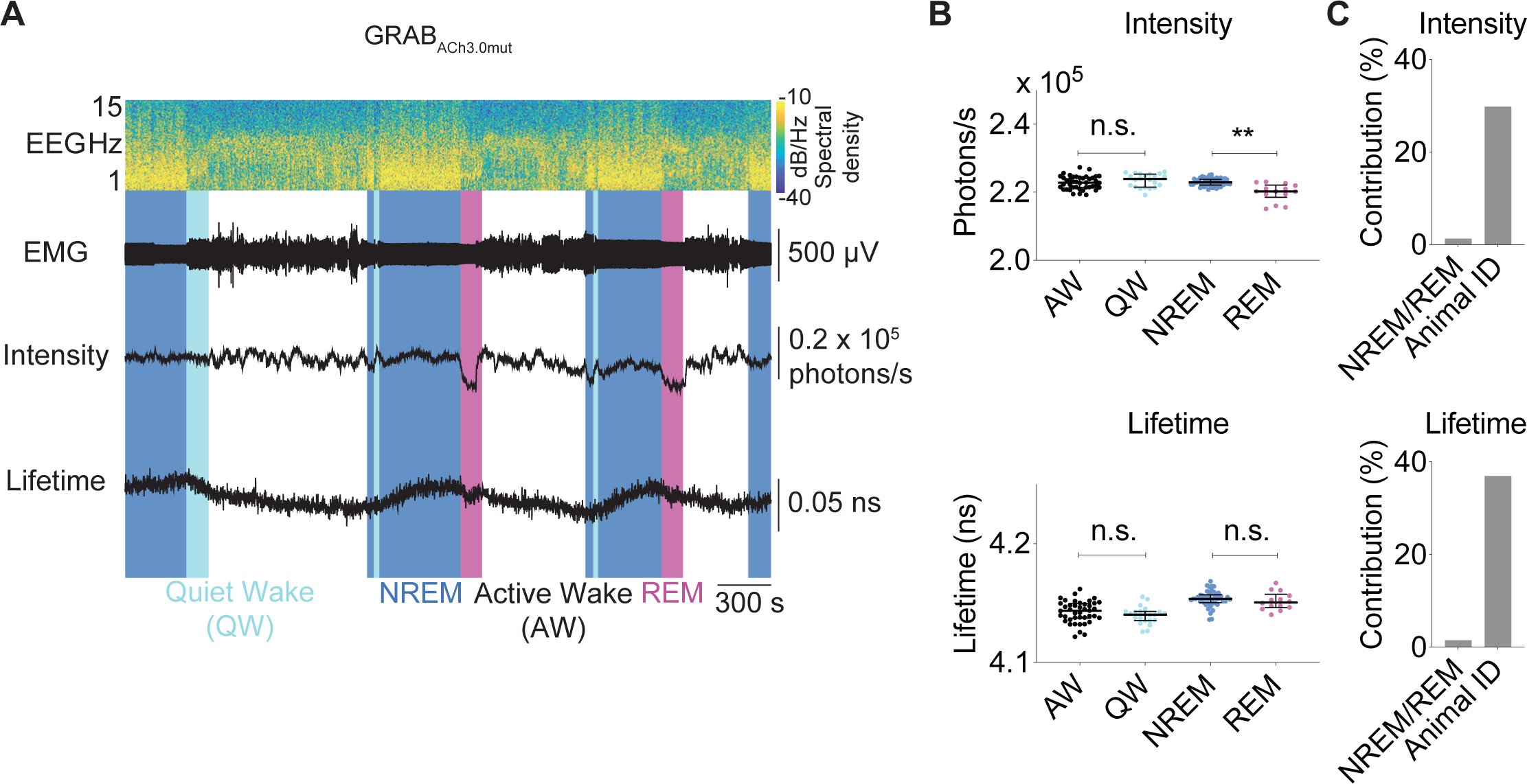
Intensity and fluorescence lifetime measurements of GRAB_ACh3.0mut_ sensor in the hippocampus across sleep-wake cycles. **(A)** Example of spectrogram of EEG recording, EMG trace, the corresponding scored sleep-wake states, along with intensity and fluorescence lifetime traces of GRAB_ACh3.0mut_ sensor from a mouse within 1 hour. Note the decrease in intensity during REM state. **(B)** Summary of intensity and fluorescence lifetime measurements of GRAB_ACh3.0mut_ sensor in different sleep-wake states. Kruskal-Wallis test with Dunn’s multiple comparison, **p < 0.01, n.s. not significant. Data are represented as median with interquartile range. **(C)** Results from two-way ANOVA analysis showing the contribution to the total variance of the measurements due to behavior states (NREM vs REM) or animal identities. Note that behavior state (NREM or REM) gives minimal contribution to the total variance of the measurements.

